# Cancer gene therapy by NF-κB-activated cancer cell-specific expression of CRISPR/Cas9 targeting to telomere

**DOI:** 10.1101/553099

**Authors:** Wei Dai, Jian Wu, Danyang Wang, Jinke Wang

## Abstract

NF-κB has been a luring target for cancer therapy due to its over activation in all tumors. In this study, we showed that a gene therapy named as NF-κB-activated gene expression (Nage) could be used to induce cancer cell death *in vitro* and *in vivo* by utilizing the NF-κB activity in cancer cells; however, it had no effect on normal cells. In this gene therapy, we constructed a NF-κB-specific promoter by fusing a NF-κB decoy sequence to a minimal promoter, which could be bound by the intracellular over activated NF-κB and thus activate the expression of downstream effector gene in a NF-κB-specific manner. In this study, we firstly demonstrated the cancer cell-specific activation of NF-κB. We then demonstrated the cancer cell specificity of Nage vector expression by introducing a Nage vector that could express a reporter gene ZsGreen in various cell lines. We next demonstrated that a Nage vector that could express CRISPR/Cas9 protein and a telomere-targeting sgRNA could be used to specifically induce death of cancer cells. We finally showed that the Cas9/sgRNA Nage vector packaged into the adeno-associated virus (AAV) could be used to inhibit the growth of xenografted tumors in mouse by intravenously injecting recombinant AAV.

## 1. Introduction

Nuclear factor-kappa B (NF-κB) is a sequence-specific DNA-binding transcription factor [1], it has been a luring target for cancer therapy due to its over activation in all cancers [2]. NF-κB is an inducible sequence-specific DNA-binding transcription factor that plays critical role in immune response [3–5]. Its five members can form hetero-- and homodimers to bind variant DNA sequences named as κB in genomes [6]. NF-κB plays its functions by regulating the expression of its target genes via binding its DNA-binding sites in cis regulatory elements [7–10]. NF-κB is found to be constitutively activated in inflammation and cancers [2, 11–14]. The active NF-κB turns on the expression of genes that keep the cell proliferation and prevent cell from apoptosis [15]. NF-κB signaling pathway is involved in the proliferation and pathogenesis of tumors via regulation of cell proliferation [16], control of apoptosis [17], promotion of angiogenesis, and stimulation of invasion and metastasis [18, 19]. Recently, NF-κB was demonstrated playing key function in the sustainment of cancer stemness [20]. Due to its always over activation in almost all cancer cells, NF-κB has been a attracting target for cancer therapy. Therefore, countless NF-κB inhibitors have been developed in past decades [10, 21, 22]; however, they rarely become clinical drugs due to their uncontrollable side effects. For example, a most promising NF-κB inhibitor, NF-κB Decoy Oligonucleotide Ointment Drug for Atopic Dermatitis (AnGes MG, Inc.), has recently failed in the phase 3 clinical trial in Japan.

A multitude of studies have demonstrated the fundamental role of NF-κB over activation in chronic inflammation and its final ending of cancers [12, 23–27]. These increasing evidence strengthens NF-κB as a luring therapeutic target of inflammation and cancer. However, many NF-κB activity inhibiting measures have been failed in clinical trial. This fact suggests that the therapeutic strategy of inhibiting NF-κB activity, directly or indirectly, is impracticable. The key underlying reason is that NF-κB is a double-edged sword [28, 29]. Despite its detrimental roles in disease as inflammation and cancer, the basic NF-κB activity is indispensable to cell normal functions such as tissue homeostasis and immunity. In other words, NF-κB-targeting strategies lack cancer cell specificity. The current uncontrollable inhibition often lead to over inhibition to its activity. To overcome this limitation, some new studies started to change strategy by focusing on inhibiting NF-κB target gene rather other NF-κB itself. For example, a D-tripeptide inhibitor, DTP3 (Ac-D-Tyr-D-Arg-D-Phe-NH2), had been developed to targetedly inhibit the NF-κB target gene GADD45β, a NF-κB-regulated antiapoptotic factor [30, 31]. This GADD45β inhibitor could disrupt the interaction of GADD45β with a JNK kinase MKK7, leading to restoring the MKK7 kinases activity and, ultimately, the JNK-mediated cell death pathway. As a result, DTP3 made potent apoptosis of multiple myeloma cells in vitro and in vivo, providing a promising candidate drug to multiple myeloma.

To develop new strategy to treat diseases by using NF-κB as a target, we recently developed a new type NF-κB inhibitor by combining NF-κB-targeting decoy and microRNA, which can sense the intracellular NF-κB activity and express an artificially designed microRNA targeting NF-κB RelA [32]. In this inhibitor, a NF-κB decoy sequence fused with a minimal promoter was used as a NF-κB responsive promoter (named as DMP) to control the expression of RelA-targeting microRNA. We found that this inhibitor can inhibit NF-κB activity in a NF-κB self-controlling manner by a feed-back loop. This strategy addressed a key problem of traditional NF-κB inhibitor, their *in vivo* application is uncontrollable. By inhibiting NF-κB activity, this inhibitor could induce apoptosis of cancer cell HepG2 but not normal liver cell HL7702 [32]. In addition, when cells were pre-treated with NF-κB inhibitor, BAY11-7082, it significantly led to the down-regulation of reporter gene, preliminarily indicating the NF-κB specificity of DMP as a NF-κB responsive promoter [32]. Therefore, we deduced that DMP was a NF-κB-specific promoter, which may be employed to develop a cancer cell-specific gene expression technique.

There is a Chinese tale about the first king of Xia Dynasty, Yu. His father, named Gun, was assigned by king Shun to control flood. He adopted to build dams to block flood. However, he failed and then killed by Shun. Then his son, Yu, was assigned by Shun to address the problem. Contrary to his father’s strategy, he used the strategy of dredging the river. He finally successfully controlled flood and became the new king. Inspired by the Yu’s strategy of controlling flood and based on our persistent studies of NF-κB [33–43], we thought that the constitutive activity of NF-κB in inflammatory and cancer cells may be utilize to treat these diseases. This strategy is contrary to the traditional inhibition of NF-κB. Based on this idea, we developed a cancer cell-specific NF-κB-activating gene expression (Nage) vector in this study. When the expression vector was transfected into tumor cells, the highly activated NF-κB could bind DMP and thus activate the expression of downstream effector gene CRISPR/Cas9. When a sgRNA targeting telomere DNA (TsgRNA) was simultaneously expressed with Cas9, the associated Cas9-TsgRNA could target to telomere DNA of cells, resulting in telomere DNA being cut and becoming shortening. The telomere DNA damage could promote cancer cell death. By compacting the TsgRNA-DMP-Cas9 expression cassette into AAV vector and intravenously injecting recombinant AAV (rAAV) to tumor-bearing mice, the Nage vector could inhibit tumor growth.

## 2. Materials and methods

### 2.1 Cell culture

293T, HepG2, HeLa, PANC-1, MDA-MB-453, RAW264.7, and Hepa1-6 cell lines were cultured in Dulbecco’s Modified Eagle Medium (DMEM) (HyClone), supplemented with 10% fetal bovine serum (FBS) (HyClone), 100 units/mL penicillin and 100 μg/mL streptomycin. A549, HT-29, SKOV-3, MRC-5, and HL7702 cell lines were cultured in Roswell Park Memorial Institute (RPMI) 1640 Medium (HyClone), supplemented with 10% FBS (HyClone), 100 units/mL penicillin and 100 μg/mL streptomycin. All cell lines were obtained from the cell resource center of Shanghai Institutes for Biological Sciences, Chinese Academy of Sciences and maintained in 5% CO_2_ at 37 °C.

### 2.2 Western blotting

Western blotting was done referring to previous studies [44]. Briefly, nuclear protein was extracted from 293T, HepG2, HeLa, PANC-1, MDA-MB-453, A549, HT-29, SKOV-3, Hepa1-6, RAW264.7, MRC-5, and HL7702 cell lines with the Nuclear and Cytoplasmic Protein Extraction Kit (Beyotime) according to the manufacturer’s instruction. The concentration of nuclear protein was determined with the BCA Protein Assay Kit (Beyotime) according to the manufacturer’s instruction. Then 10 μg of nuclear protein extracts were separated by 10% SDS-polyacrylamide gel electrophoresis and transferred to PVDF membrane (Millipore). After blocking with QuickBlockTM Blocking Buffer for Western Blot (Beyotime), the membranes were incubated overnight at 4 °C with the primary antibodies NF-κB p65 Mouse mAb (Cell Signaling Technology, diluted 1:1000) or TBP Rabbit mAb (Cell Signaling Technology, diluted 1:1000). After washing three times with 1 M TBS (pH 7.6) containing 0.1% (v/v) Tween 20, the membranes were then incubated 1 h at room temperature with second antibodies IRDye^®^ 800CW Goat anti-Mouse IgG (LI-COR, diluted 1:10000) or IRDye^®^ 800CW Goat anti-Rabbit IgG (LI-COR, diluted 1:10000). Finally, membranes were washed three times with 1 M TBS (pH 7.6) containing 0.1% (v/v) Tween 20, and scanned with Infrared Imaging System (LI-COR), and the protein expression values of NF-κB were normalized with that of internal reference TBP using Odyssey V3.0 software.

### 2.3 Vector construction

Two complementary oligonucleotides (DMP-F and DMP-R; Table S1) consisting of five NF-κB binding sites (GGGAATTTCC GGGGACTTTCC GGGAATTTCC GGGGACTTTCC GGGAATTTCC) regarded as NF-κB decoy and a minimal promoter sequence (TAGAGGGTATATA) were annealed, extended and amplified with primers DMP XS-F and DMP EB-R (Table S1) to generate DMP-1 fragment, then subcloned into T-clone vector pMD19-T (TaKaRa) to generate pT-DMP-1. Linear ZsGreen fragment was amplified from plasmid pLVX-shRNA-ZsGreen (Addgene) with primers ZsGreen E-F and ZsGreen B-R (Table S1), then cloned into pT-DMP-1 using the cutting sites of EcoRI and BamHI enzymes (ThermoFisher Scientific) to generate pDMP-ZsGreen.

Above NF-κB decoy and minimal promoter sequence were annealed, extended and amplified with primers DMP ENB-F and DMP XHS-R (Table S2) to generate DMP-2 fragment, then subcloned into T-clone vector pMD19-T (TaKaRa) to generate pT-DMP-2. Two complementary oligonucleotides (TsgRNA-F and TsgRNA-R; Table S2) with a 20-bp targeting telomere sequences (TAGGGTTAGGGTTAGGGTTA) were annealed and cloned into px458 (Addgene) using the digestion sites of BbsI (NEB) to generate pTsgRNA-Cas9. Linear TsgRNA fragment was amplified from pTsgRNA-Cas9 with primers U6 NcoI-F and U6 BamHI-R (Table S2), then cloned into pT-DMP-2 using the cutting sites of NcoI and BamHI enzymes (ThermoFisher Scientific) to generate pT-TsgRNA-DMP. Linear Cas9-EGFP fragment was amplified from px458 (Addgene) with primers Cas9 SalI-F and Cas9 HindIII-R (Table S2), then cloned into pT-TsgRNA-DMP using the cutting sites of SalI and HindIII enzymes (ThermoFisher Scientific) to generate pTsgRNA-DMP-Cas9-EGFP.

TsgRNA-DMP-Cas9 fragment was cut out from pTsgRNA-DMP-Cas9-EGFP, then cloned into pAAV-MCS vector (Stratagene) using the cutting sites of EcoRI and HindIII enzymes (ThermoFisher Scientific) to generate pAAV-TsgRNA-DMP-Cas9.

### 2.4 Cell transfection

In order to investigate the cancer cell-specificity of Nage vector, 293T, HepG2, HeLa, PANC-1, MDA-MB-453, A549, HT-29, SKOV-3, Hepa1-6, RAW264.7, MRC-5, and HL7702 cell lines were seeded into 24-well plates at a density of 0.5×105 cells/well and cultivated overnight. Then cells were transfected 500 ng of pDMP-ZsGreen or pEGFP-C1 (Addgene) using Lipofectamine^®^ 2000 Reagent (ThermoFisher Scientific) according to the following protocol. For each transfection, cells were cultured with 500 μL of Opti-MEM^®^ Medium (ThermoFisher Scientific) at 37 °C for 30 min. Two stock solutions were prepared for each transfection, one was 50 μL of Opti-MEM, 500 ng of total DNA, another was 50 μL of Opti-MEM, 2 μL of Lipofectamine 2000, respectively. The two solutions were vortexed and incubated for 5 min at room temperature, respectively. Then, the Opti-MEM/Lipofectamine solution was added to the Opti-MEM/DNA solution in dropwise, vortexed, and incubated for 20 min at room temperature before being added to each well. After incubated for 5 h, the medium of each well was replaced with 500 μL of fresh DMEM or RPMI 1640 medium containing 10% FBS. The cells were incubated in 5% CO_2_ at 37 °C for another 24 h. All cells were then observed and photographed with a fluorescence microscope (IX51; Olympus) at the constant magnification of 200×. The fluorescence intensity of cells were quantified with a flow cytometry (Calibur; BD), and the mean fluorescence intensity (MFI) was calculated by BD software of flow cytometry.

In order to investigate whether the Nage vector could be used to kill cancer cells, 293T, HepG2, HeLa, PANC-1, MDA-MB-453, A549, HT-29, SKOV-3, Hepa1-6, RAW264.7, MRC-5, and HL7702 cells were seeded into 24-well plates and transfected with 500 ng of pTsgRNA-DMP-Cas9-EGFP or pEGFP-C1 (Addgene) using Lipofectamine^®^ 2000 Reagent (ThermoFisher Scientific) according to the above protocol. After transfection, cells were cultivated with the fresh medium containing 10% FBS for 24 h. Then cells were washed twice with PBS and stained with 100 μg/mL acridine orange (Solarbio) for 10 min at room temperature. All cells were observed and photographed with fluorescent fluorescence microscope (IX51; Olympus) at the constant magnification of 200×.

### 2.5 Cytotoxicity assay

In order to detect cell viability, 293T, HepG2, HeLa, PANC-1, MDA-MB-453, A549, HT-29, SKOV-3, Hepa1-6, RAW264.7, MRC-5, and HL7702 cell lines were seeded into 96-well plates at a density of 1×10^4^ cells/well and cultivated overnight. Then cells were transfected with 100 ng of pTsgRNA-DMP-Cas9-EGFP or pEGFP-C1 (Addgene) using Lipofectamine^®^ 2000 Reagent (ThermoFisher Scientific) according to the manufacturer’s instructions. After transfection, cells were cultivated with the fresh medium containing 10% FBS (100 μL/well) for 5 h, then added with 10 μL of alamar blue (Yeasen) per well. Cells were then cultivated for 24 h. The fluorescence intensity of each well was measured with a microplate reader (Synergy HT; BioTek) by using an excitation wavelength at 530 nm and an emission wavelength at 590 nm. The cell survival rate of treated cells was calculated as the percentage of fluorescence intensity of treated cells wells to that of control wells.

### 2.6 Telomere length measurement

The telomere measurement by quantitative PCR was performed as previous study [45]. Briefly, the genomic DNA (gDNA) was extracted from 293T, HepG2, HeLa, PANC-1, MDA-MB-453, A549, HT-29, SKOV-3, Hepa1-6, RAW264.7, MRC-5, and HL7702 cell lines after transfection with pTsgRNA-DMP-Cas9-EGFP 24 h using the TIANamp Genomic DNA Kit (Tiangen) according to the manufacturer’s instructions. The quantitative telomere length was analyzed by qPCR with primers Tel qF and Tel qR (Table S3). The single copy internal control of β-globin gene was analyzed by qPCR with primers β-globin qF and β-globin qR (Table S3). The qPCR reaction (20 μL) contained 10 μL of SYBR Green Real-time PCR Master Mix (2×) (Roche), 0.25 μM primers, and 10 ng of gDNA. For telomere qPCR, the program was as follows: 95 °C for 10 min, 22 cycles of 95 °C for 15 s and 54 °C for 2 min. For β-globin gene qPCR, the program was as follows: 95 °C for 10 min, 30 cycles of 95 °C for 15 s and 58 °C for 1 min. A melting curve was conducted to monitor the PCR amplification. The qPCR programs were run on a StepOne Plus real-time PCR machine (Applied Biosystems). Each qPCR detection was performed in at least three technical replicates. Melting curve analysis revealed a single PCR product. Data analysis was performed using the Applied Biosystems StepOne software v2.3, and Ct values of telomere (T) were normalized with that of single copy gene β-globin (S). The relative telomere length was calculated as T/S ratio according to the equation: RQ = 2^ΔCt^.

### 2.7 rAAV preparation

293T cells were plated in 75-cm^2^ flask at a density of 5×10^6^ cells/flask and cultivated overnight. Then cells were transfected with pAAV-TsgRNA-DMP-Cas9 and pAAV-MCS using Lipofectamine^®^ 2000 Reagent (ThermoFisher Scientific) according to the following protocol. Before transfection, 293T cells were cultured with 15 mL of Opti-MEM^®^ Medium (ThermoFisher Scientific) at 37 °C for 30 min. Two stock solutions were prepared for each transfection. One was 1 mL of Opti-MEM plus 4 μg of pAAV-TsgRNA-DMP-Cas9 or pAAV-MCS, 4 μg of pAAV-RC (Stratagene), and 4 μg of pHelper (Stratagene). The other was 1 mL of Opti-MEM plus 45 μL of Lipofectamine^®^ 2000. The two solutions were then vortexed and incubated at room temperature for 5 min, respectively. Then the Opti-MEM/Lipofectamine solution was added to the Opti-MEM/DNA solution in dropwise. The mixture was vortexed and incubated at room temperature for 20 min before added to cells in a flask. After incubated for 5 h, the medium of each flask was replaced with 15 mL of fresh DMEM medium containing 10% FBS. Cells were cultivated for 72 h.

In order to prepare viral lysates, the transfected 293T cells were scraped off with a cell scraper and transferred the cell suspension into 50-mL screw-cap conical tubes. The cell suspension was subjected three rounds of freeze-thaw-vortex cycles to release rAAV from cells as follows: froze cells at −70 °C for 2 h, thawed cells in a 37 °C water bath, and vortexed vigorously. Then cell debris was removed by centrifugation at 500 g at room temperature for 10 min and the supernatant was collected as the initial rAAV lysate. The supernatant was pooled into a 50-mL conical tube, added 1/10 volume of chloroform and incubated at 37°C for 1 h with rotation. Solid sodium chloride was then added to the mixture at a final concentration of 1 M. The supernatant was collected by centrifuging at 10,000 g at 4 °C for 15 min and transferred into a new 50-mL conical tube. The supernatant was then added with PEG8000 (Sigma) at a final concentration of 10%. The mixture was incubated on ice for 1 h. The mixture was then centrifuged at 10,000 g at room temperature for 15 min to remove the supernatant. The pellet was resuspended with 5 mL of PBS buffer, which was used as intermediate rAAV product. In order to concentrate and purify the rAAV, the intermediate rAAV product was added with DNase and RNase at a final concentration of 1 μg/mL and incubated at room temperature for 30 min. Then the intermediate rAAV product was added with the same volume of chloroform and mixed by rotation. The mixture was centrifuged at 10,000 g at 4 °C for 10 min to collect the supernatant, which was used as the final rAAV product. Two final rAAV products were prepared, named as rAAV-TsgRNA-DMP-Cas9 and rAAV-MCS.

### 2.8 rAAV titration

The titration of rAAV was analyzed by quantitative PCR (qPCR). The qPCR amplification of rAAV-TsgRNA-DMP-Cas9 and rAAV-MCS used the primers DMP qF/R and MCS qF/R, respectively (Table S3). A purified TsgRNA fragment with known molecule copy number was used as standard sample to establish a standard curve, which was 10-fold serially diluted at the concentrations ranging from 1:10 to 1:10^4^. The qPCR amplification of TsgRNA fragment used the primers U6 F and U6 R (Table S3). The PCR reaction (20 μL) consisted of 10 μL of SYBR Green Real-time PCR Master Mix (2×) (Roche), 0.25 μM primers, and 2 μL of purified rAAV or TsgRNA fragment. The PCR programs were as follows: 95 °C for 10 min and 45 cycles of 95 °C for 15 s and 60 °C for 1 min. A melting curve was conducted to monitor the PCR amplification. The qPCR programs were run on a StepOne Plus real-time PCR machine (Applied Biosystems). Each qPCR detection was performed in at least three technical replicates. Melting curve analysis revealed a single PCR product. Data analysis was performed using the Applied Biosystems StepOne software v2.3, the concentration of virus genome (vg) was determined according to the standard curve.

### 2.9 rAAV evaluation

In order to evaluate the prepared rAAVs, HepG2, Hepa1-6, MRC-5, and HL7702 cells were seeded into 24-well plates at a density of 0.5×10^5^ cells/well and cultivated overnight. Cells were then transfected with the viruses rAAV-TsgRNA-DMP-Cas9 and rAAV-MCS at the dose of 1×10^2^ vg/cell, respectively. The transfected cells were cultivated for 24 h. Cells were then stained with acridine orange as described above. All cells were observed and photographed with a fluorescence microscope (IX51; Olympus) at the constant magnification of 200×.

### 2.10 Animal experiment

Four-week-old female nude mice (BALB/c-Foxn1^nu^) purchased from the Model Animal Research Center of Nanjing (Nanjing, China) were used as the experimental animals. Each mouse weighed 18-22 g and was maintained under specific pathogen-free conditions. All animal experiments in this study followed the guidelines and ethics of the Animal Care and Use Committee of Southeast University (Nanjing, China).

In the first animal experiment, the nude mice was subcutaneously transplanted with the mixture of 1×10^7^ Hepa1-6 cells and 1×10^9^ vg rAAV-MCS or rAAV-TsgRNA-DMP-Cas9. On each mouse, the mixture of Hepa1-6 and rAAV was transplanted on both sides of abdomen. After two weeks, all mice were euthanized and photographed. Tumor size was measured with a precision caliper. Tumor volume was calculated with the formula V = (D*d*^2^)/2, in which *D* is the major tumor axis and *d* is the minor tumor axis.

In the second animal experiment, the nude mice was subcutaneously transplanted with 1×10^7^ Hepa1-6 cells on both sides of abdomen to produce the tumor-bearing mice. After one week, the tumor-bearing mice were randomly divided into two groups and intravenously injected with 1×10^9^ vg of rAAV-MCS or rAAV-TsgRNA-DMP-Cas9, respectively. After one week, all mice were euthanized and photographed. Tumor size was measured and calculated as described above.

### 2.11 Virus detection

In order to detect the presence of virus in tissues with qPCR, the heart, liver, spleen, lung, kidney, and tumor tissues were collected from mice in the second animal experiment. The genomic DNA (gDNA) was extracted from above tissues using the TIANamp Genomic DNA Kit (TIANGEN) according to the manufacturer’s instructions. The rAAV DNA abundance was analyzed by qPCR with primers MCS qF and MCS qR (Table S3). The internal control of *GAPDH* gene was analyzed by qPCR with primers GAPDH qF1 and GAPDH qR1 (Table S3). The qPCR reaction (20 μL) contained 10 μL of SYBR Green Real-time PCR Master Mix (2×) (Roche), 0.25 μM primers, and 10 ng of gDNA. The qPCR program was as follows: 95 °C for 10 min, 45 cycles of 95 °C for 15 s and 60 °C for 1 min. A melting curve was conducted to monitor the PCR amplification. The qPCR programs were run on a StepOne Plus real-time PCR machine (Applied Biosystems). Each qPCR detection was performed in at least three technical replicates. Melting curve analysis revealed a single PCR product. Data analysis was performed using the Applied Biosystems StepOne software v2.3, and Ct values were normalized with that of *GAPDH*. The relative abundance of viral DNA was calculated as relative quantity (RQ) according to the equation: RQ = 2^−ΔΔCt^.

### 2.12 RT-PCR

In order to detect the expression of RelA and Cas9 in tissues with qPCR, the heart, liver, spleen, lung, kidney, and tumor tissues were collected from mice in the second animal experiment. Total RNA was extracted from above tissues with Trizol^®^ reagent (Invitrogen), then 500 ng of total RNA was reversely transcribed into cDNA using PrimeScriptTM RT Master Mix (TaKaRa) according to the manufacturer’s instructions in a total volume of 10 μL. Then RelA and Cas9 mRNA expression levels were analyzed by qPCR with primers RelA qF/R and Cas9 qF/R (Table S3). The internal control of *GAPDH* gene was analyzed by qPCR with primers GAPDH qF2 and GAPDH qR2 (Table S3). The qPCR reaction (20 μL) contained 10 μL of 2× SYBR Green Real-time PCR Master Mix (Roche), 0.25 μM primers, and 2 μL of cDNA. The qPCR program was as follows: 95 °C for 10 min, 45 cycles of 95 °C for 15 s and 60 °C for 1 min. A melting curve was conducted to monitor the PCR amplification. The qPCR programs were run on a StepOne Plus real-time PCR machine (Applied Biosystems). Each qPCR detection was performed in at least three technical replicates. Melting curve analysis revealed a single PCR product. Data analysis was performed using the Applied Biosystems StepOne software v2.3, and Ct values were normalized with that of *GAPDH*. The relative RelA and Cas9 mRNA expression level was calculated as relative quantity (RQ) according to the equation: RQ = 2^−ΔΔCt^.

### 2.13 Statistics

Data were expressed as means values ± standard deviation (SD). The statistical significance was analyzed by a two-tailed unpaired Student’s *t*-test. Differences at *p* < 0.05 were considered statistical significance.

## 3. Results

### 3.1 Evaluation of NF-κB activity in different cell lines

Because the intracellular NF-κB activity is critical to Nage vector, we firstly detected the NF-κB activity in twelve cell lines, including seven human cancer representing different cancers (HepG2, A549, PANC-1, MDA-MB-453, HT-29, HeLa, and SKOV-3), two mouse cancer cell lines (Hepa1-6 and RAW264.7), and two normal human cell lines (MRC-5 and HL7702). We also detected the NF-κB activity in an engineered normal human cell lines (293T), which was widely used to transfect exogenous genes with high efficiency. We expected to known the cancer cell-specific over activation of NF-κB. Because the main functional NF-κB protein is the heterodimer of RelA-p50 and the activated NF-κB protein locates in nucleus, we detected the abundance of RelA protein in nuclear extracts with Western blot by using a anti-RelA antibody. Another reason of detecting RelA is that only this protein has transcriptional activation domain in RelA-p50 dimer, which exerts the NF-κB’s function of gene transcriptional activation. The results indicated that a single 64-kDa band of NF-κB protein was detected in 293T and all other tumor cells, including HepG2, HeLa, PANC-1, MDA-MB-453, A549, HT-29, SKOV-3, Hepa1-6, and RAW264.7 cells (Figure 1a). However, no NF-κB protein was found in two normal cells (MRC-5 and HL7702) (Figure 1a). In this Western blot detection, the nuclear protein TBP was simultaneously detected as a loading control (a single 37-kDa band) (Figure 1a). In contrast to HL7702, all cancer cell lines and 293T had significantly high NF-κB activity (Figure 1b). However, the relative NF-κB activity is not identical among different cell lines, showing cell type-dependent NF-κB activation level. It was found there was the highest relative NF-κB activity in 293T cell. These results indicated that NF-κB was only constitutively over activated in all detected cancer cells.

**Figure 1.**
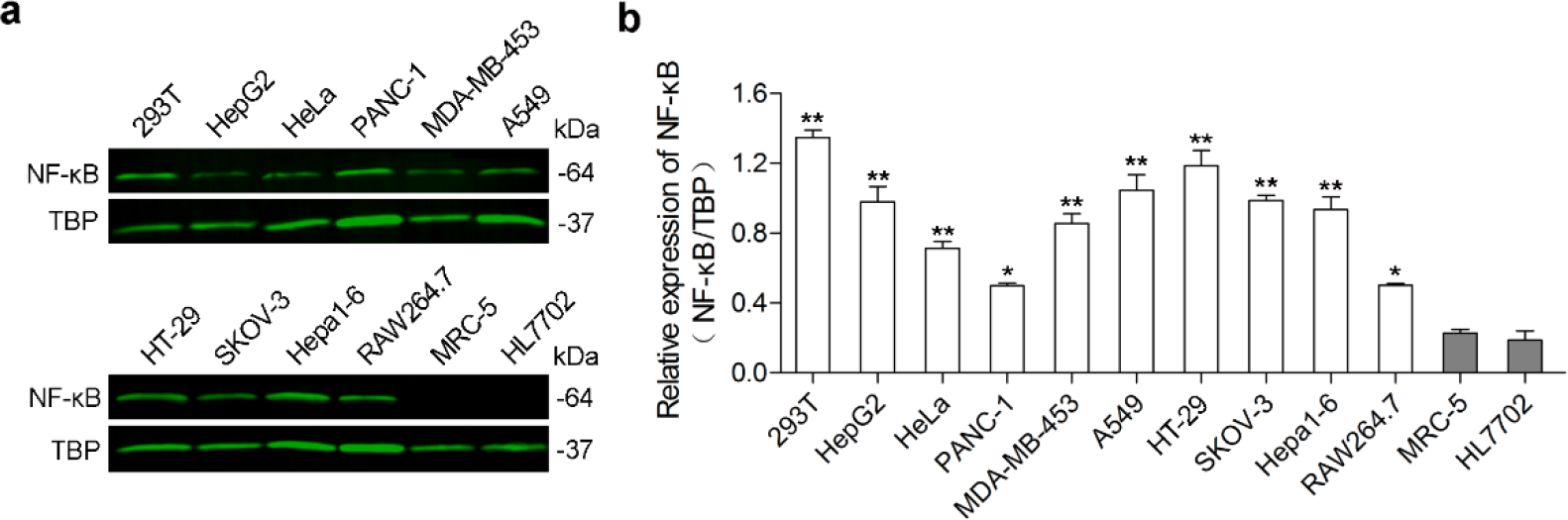
Activation of NF-κB protein in various cell lines. Nuclear protein was extracted from cells and detected with western blot by using a anti-RelA/p65 antibody. a, Western blot detection of NF-κB p65 and TBP protein abundance in nuclear protein. b, Relative NF-κB activity in cells. TBP was used as a loading control of nuclear protein to normalize the abundance of NF-κB. The statistical significance was checked between the normalized NF-κB activity in all cancer cell lines and that of HL7702 cell. *, *p* ≤ 0.05; **, *p* ≤ 0.01.

### 3.2 Activation of reporter gene expression in cells by Nage vector

In this study, we expect to induce cancer cell death by transfecting a Nage vector. In this vector, we used a NF-κB-specific promoter to drive expression of a downstream effector gene that can induce cell death. The NF-κB-specific promoter was a DNA sequence formed by fusing a NF-κB decoy sequence with a minimal promoter [32], which was therefore named as DMP, referring to decoy minimal promoter. To investigate the cancer cell-specific gene activation of Nage vector, we constructed a plasmid vector named DMP-ZsGreen by cloning ZsGreen gene downstream of DMP (Figure 2a). We conceived that the expression of ZsGreen protein could be activated by NF-κB activity in cancer cells but not in normal cells without NF-κB activity (Figure 2a). We thus transfected all cell lines detected above with the DMP-ZsGreen plasmid. The results indicated that ZsGreen was successfully expressed in 293T and all cancer cells (Figure 2b). By quantifying the fluorescence intensity of as many as 10,000 cells with flow cytometry, the MFI was measured to be 15% (293T), 8.3% (HepG2), 8.7% (HeLa), 10% (PANC-1), 1.8% (MDA-MB-453), 8.3% (A549), 2.2% (HT-29), 6.2% (SKOV-3), 8.3% (Hepa1-6), and 1.4% (RAW264.7), respectively (Figure 2b). In contrast, ZsGreen was not expressed in normal cells including MRC-5 and HL7702 when transfected with the same plasmid (Figure 2b). As a blank control, all cells only transfected lipofectin showed no fluorescence (Figure S1).

**Figure 2.**
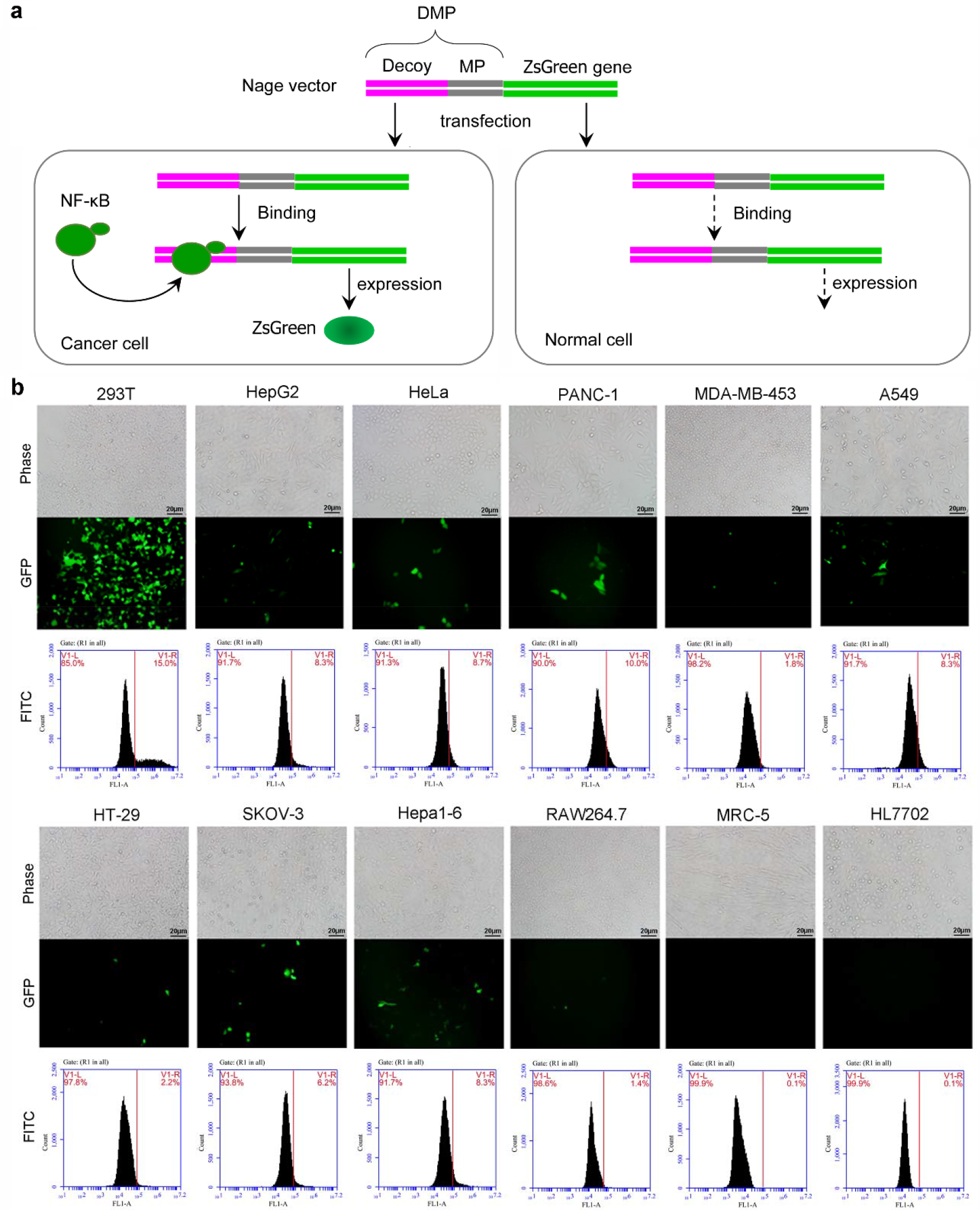
Expression of ZsGreen with Nage vector in various cell lines. a, Schematic of Nage vector to activate ZsGreen expression. MP, minimal promoter; DMP, decoy minimal promoter; Nage, NF-κB activating gene expression. b, Representative cell images under the bright field (Phase) and fluorescence (GFP). Cells were transfected by the Nage vector expressing ZsGreen.

In this transfection, it was found that 293T cell showed the highest ZsGreen expression, in agreement with the highest NF-κB activity in this cell (Figure 1). However, in cancer cells, there is no correlation between ZsGreen expression level and NF-κB activity. For example, in all cancer cells, HT29 cell had the highest NF-κB activity but the almost lowest ZsGreen expression; however, PANC1 cell had the lowest NF-κB activity but the highest ZsGreen expression. This phenomenon may be related with the transfection efficiency of different cells. Anyway, the Nage vector was activated to express ZsGreen in all NF-κB over activated cells, including 293T and all cancer cells; however, it was not activated in all normal cells, indicating the cancer cell-specific expression of Nage vector.

As a transfection control, we also simultaneously transfected all cells with a plasmid C1-EGFP, in which the EGFP expression was driven by CMV promoter. This promoter is a widely used natural mammalian promoter with the highest transcriptional activation activity [42]. The results demonstrated that all cell lines showed EGFP expression, including two normal cell lines (MRC-5 and HL7702), indicating the cancer cell non-specificity of this powerful promoter. This result is in agreement with our previous study, in which the CMV promoter and DMP were compared by only transfecting HepG2 and HL7702 [32]. Our previous study revealed that the transcriptional activation activity of CMV promoter also had close relationship with NF-κB activity because it contains four canonical NF-κB binding sites [42]; however, this study verified that it is not a NF-κB-specific promoter. Therefore, it could drive EGFP expression in two normal cell lines that had no NF-κB activity. Finally, by comparing the MFI of cells other than two normal cells transfected by CMV-EGFP and DMP-ZsGreen, we noticed that there was a linear correlation (Figure S2), indicating that the above described no correlation between the ZsGreen expression level and the NF-κB activity resulted from the different transfection efficiency of different cells.

### 3.3 Promotion of cancer cell death *in vitro* by expressing Cas9 with a plasmid Nage vector

After validating the cancer cell-specific exogenous gene expression by Nage vector, we expected to known whether the Nage vector could be used to promote cancer cell death. Because the CRISPR/Cas9 was widely used to introduce double-strand break (DSB) to genomes of living cells and the DNA damage can induce cell apoptosis, we thus selected Cas9 as an effector gene in Nage vector. Because previous study revealed that a telomere-targeting sgRNA (TsgRNA) could guide a EGFP-fused dead Cas9 (dCas9) to an array of telomere positions for imaging telomeres in cells [46], we deduced that the Nage vector-expressed Cas9 protein could be guided to telomeres of cancer cells by a co-expressed TsgRNA, where Cas9 should be produce DSBs to telomere DNA and thus promote cancer cell death (Figure 3a). However, in normal cells without NF-κB activity, the Cas9 protein can not be expressed; accordingly, the Nage vector transfection should exert no effect on normal cells (Figure 3a). Therefore, we constructed a new plasmid Nage vector, TsgRNA-DMP-Cas9-EGFP, in which the expression of TsgRNA was under the control of U6 promoter but the expression of Cas9-EGFP was under the control of DMP (Figure 3a).

**Figure 3.**
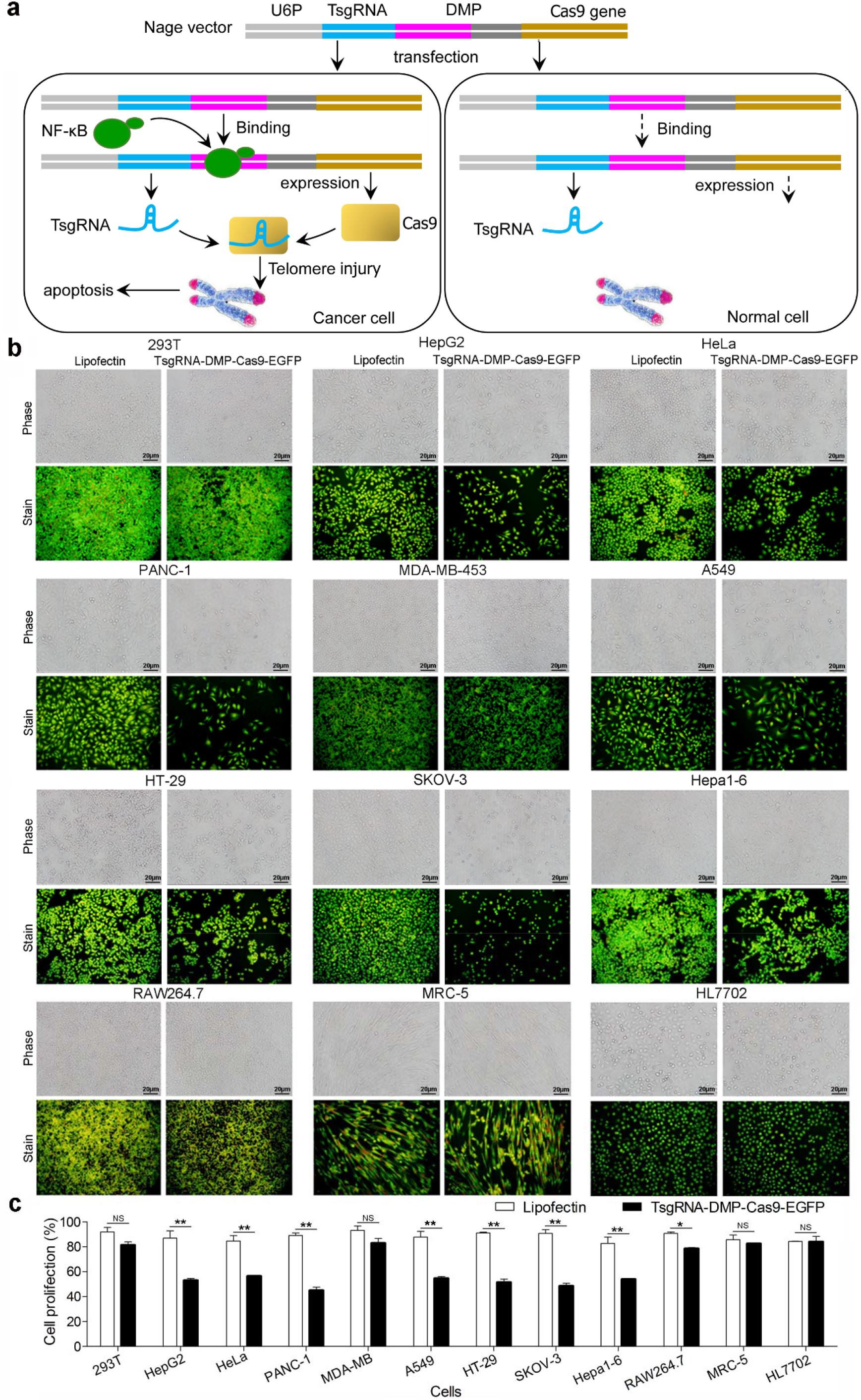
Expression of Cas9-TsgRNA with Nage vector in various cell lines. a, Schematic of killing tumor cells with Nage vector. TsgRNA, telomere-targeting sgRNA; DMP, decoy minimal promoter; U6P, U6 promoter. b, Representative cell images under the bright field (Phase) and fluorescence (Stain). Cells were transfected by the Nage vector expressing Cas9 and TsgRNA. Cells were stained with acridine orange. c, Cell viability detected by alamar blue assay. MDA-MB, MDA-MB-453; RAW, RAW264.7.

We then transfected all cells with the new plasmid Nage vector, TsgRNA-DMP-Cas9-EGFP. The results indicated that the EGFP protein was expressed in all cancer cells with different levels (Figure S3), suggesting that Cas9 protein was successfully expressed in all cancer cells by Nage vector. At the same time, as expected, the transfection with this Nage vector induced the visible death of all cancer cells (Figure 3b). In contrast, the two normal cells (MRC-5 and HL7702) showed no both EGFP expression (Figure S3) and cell death (Figure 3b) when transfected with the same Nage vector. The quantitative measurement of cell viability with alamar blue also indicated that the Nage vector transfection led to significant decrease of cell viability of all cancer cells except MDA-MB-453. In above transfection, we also simultaneously transfected all cells with pC1-EGFP (also named as pCMV-EGFP) and lipofectin and detected their effect on cell viability with alamar blue assay. The results indicated that the two transfections had no effect on cell viability (Figure S3). These results demonstrated that the Nage vector could be used to effectively and specifically induce cancer cell death by expressing Cas9 and TsgRNA. It was interesting that the TsgRNA-DMP-Cas9-EGFP transfection exerted no significant effect on 293T cells (Figure 3) despite the EGFP was most highly expressed in this cell (Figure S3).

To verify that the telomere was damaged by the Nage vector-expressed TsgRNA-Cas9, we subsequently detected the telomere length of Nage vector-transfected cells by using a qPCR method. The results indicated the telomere of all cancer cells became significant shortened with the transfection of Nage vector expressing TsgRNA-Cas9 (Figure 4). However, the telomere of two normal cells did not shorten despite transfection with Nage vector (Figure 4). Unexpectedly, the telomere of 293T was also not shortened (Figure 4). These results demonstrated that the telomere of all cancer cells could be specifically damaged by the cutting of TsgRNA-Cas9 that was expressed by the Nage vector. The death of cancer cells was induced by the telomeric DNA damage resulted from TsgRNA-Cas9 cutting.

**Figure 4.**
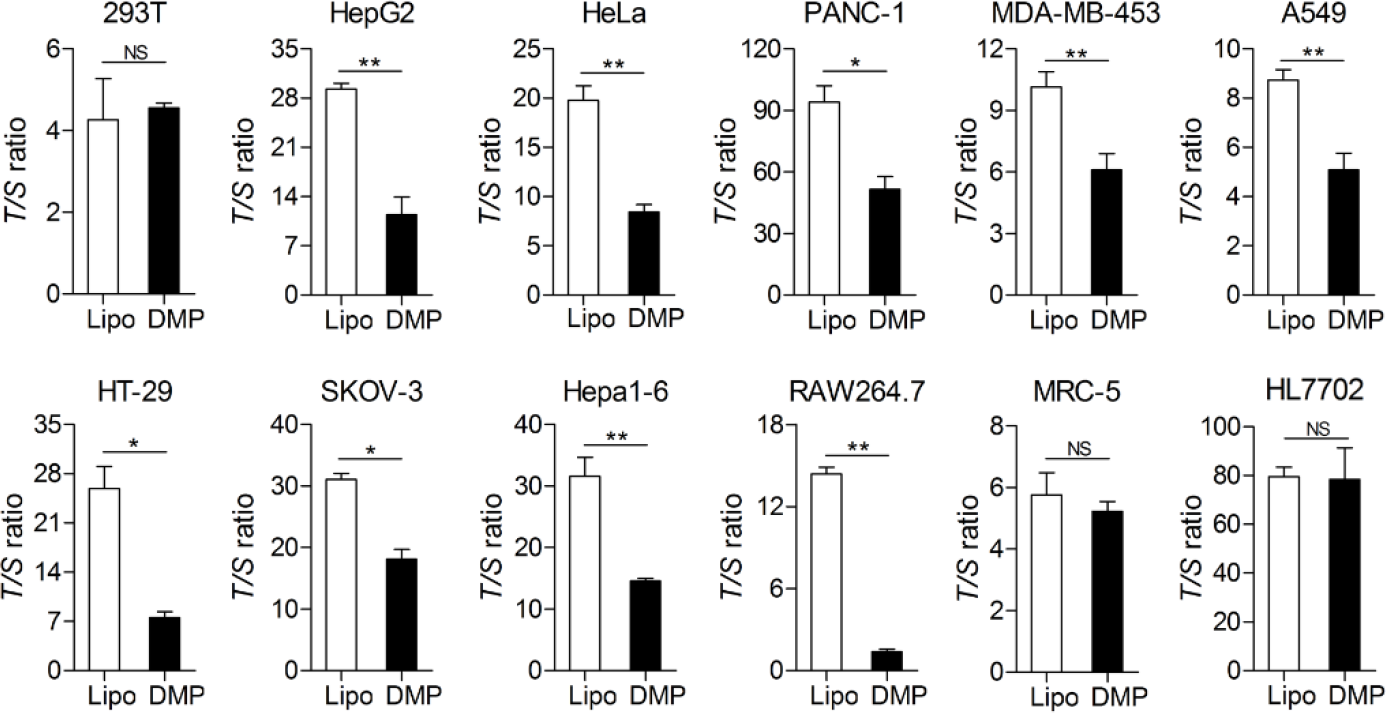
Quantitative telomere length measurement with qPCR. Cells were transfected by the Nage vector expressing Cas9 and TsgRNA. The telomere length was detected with a qPCR method. Lipo, Lipofectin; DMP, TsgRNA-DMP-Cas9-EGFP; T, telomere; S, single copy gene β-globin.

### 3.4 Promotion of cancer cell death *in vitro* by expressing Cas9 with an AAV Nage vector

To explore the *in vivo* application of the Cas9-expressing Nage vector, the whole sequence of U6 promoter, TsgRNA, DMP, and Cas9 in plasmid TsgRNA-DMP-Cas9-EGFP was cloned into an AAV vector to construct rAAV-TsgRNA-DMP-Cas9. Then, the effect of rAAV-TsgRNA-DMP-Cas9 to promote cancer cell death was evaluated by transfecting four cell lines including HepG2, Hapa1-6, MRC-5, and HL7702. The results indicated that rAAV-TsgRNA-DMP-Cas9 induced significant cell death in two cancer cells (HepG2 and Hepa1-6) (Figure 5a, b). However, it exerted no effects on two normal cells (MRC-5 and HL7702) (Figure 5c, d). As a negative control, we also simultaneously transfected the four cell lines with rAAV-MCS that was cloned no TsgRNA-DMP-Cas9 sequences (Figure 5). The results indicated that the growth of all four cell lines was not affected by the control rAAV. These results verified the effect of rAAV-TsgRNA-DMP-Cas9.

**Figure 5.**
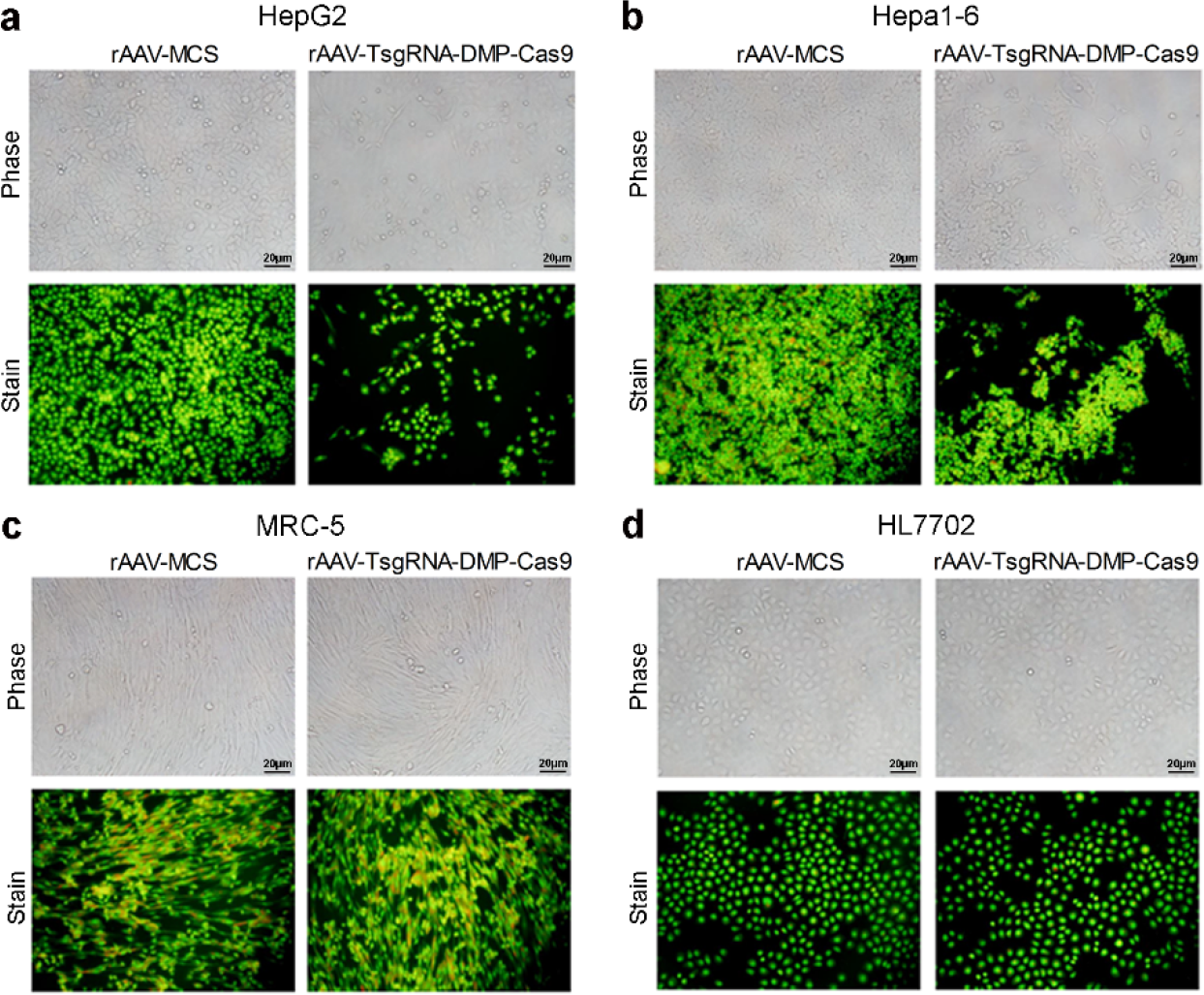
Evaluation of rAAV. Cells were transfected by the recombinant AAV packaged with the Nage vector expressing Cas9 and TsgRNA (rAAV-TsgRNA-DMP-Cas9). Cells were stained with acridine orange. a-d, Representative cell images under the bright field (Phase) and fluorescence (stain) of four cell lines, HepG2, Hepa1-6, MRC-5, and HL7702 cells, respectively.

### 3.5 Inhibition of tumor growth *in vivo* by expressing Cas9 with AAV Nage vector

To explore whether the Nage vector packaged in AAV viron could inhibit tumor growth *in vivo*, we next performed animal experiments. In the first animal experiment, the Hepa1-6 cells were firstly mixed with the control virus rAAV-MCS and rAAV-TsgRNA-DMP-Cas9 *in vitro*, respectively. Then the mixtures were subcutaneously transplanted at two sides of abdomen of nude mice at the same time. After two weeks, all nude mice transplanted with the Hepa1-6/rAAV-MCS mixture appeared obvious large tumors on both sides and tumor mass hyperemia and swelling (Figure 6a; Figure S4a). The mean tumor size of left and right side was up to 190 mm3 and 207.75 mm3, respectively (Figure 6b). However, the tumor size of nude mice transplanted with the Hepa1-6/rAAV-TsgRNA-DMP-Cas9 mixture was significantly smaller than that of control groups (Figure 6a). The mean tumor size of left and right side was only 12.58 mm3 and 15.96 mm3, respectively (Figure 6b). We noticed that many tumors completely did not grow out in mice transplanted with the Hepa1-6/rAAV-TsgRNA-DMP-Cas9 mixture (Figure 6a; Figure S4b). However, all tumors grow out in mice transplanted with the Hepa1-6/rAAV-MCS mixture. These results demonstrated the strong tumor inhibition function of rAAV-TsgRNA-DMP-Cas9 *in vivo.*

**Figure 6.**
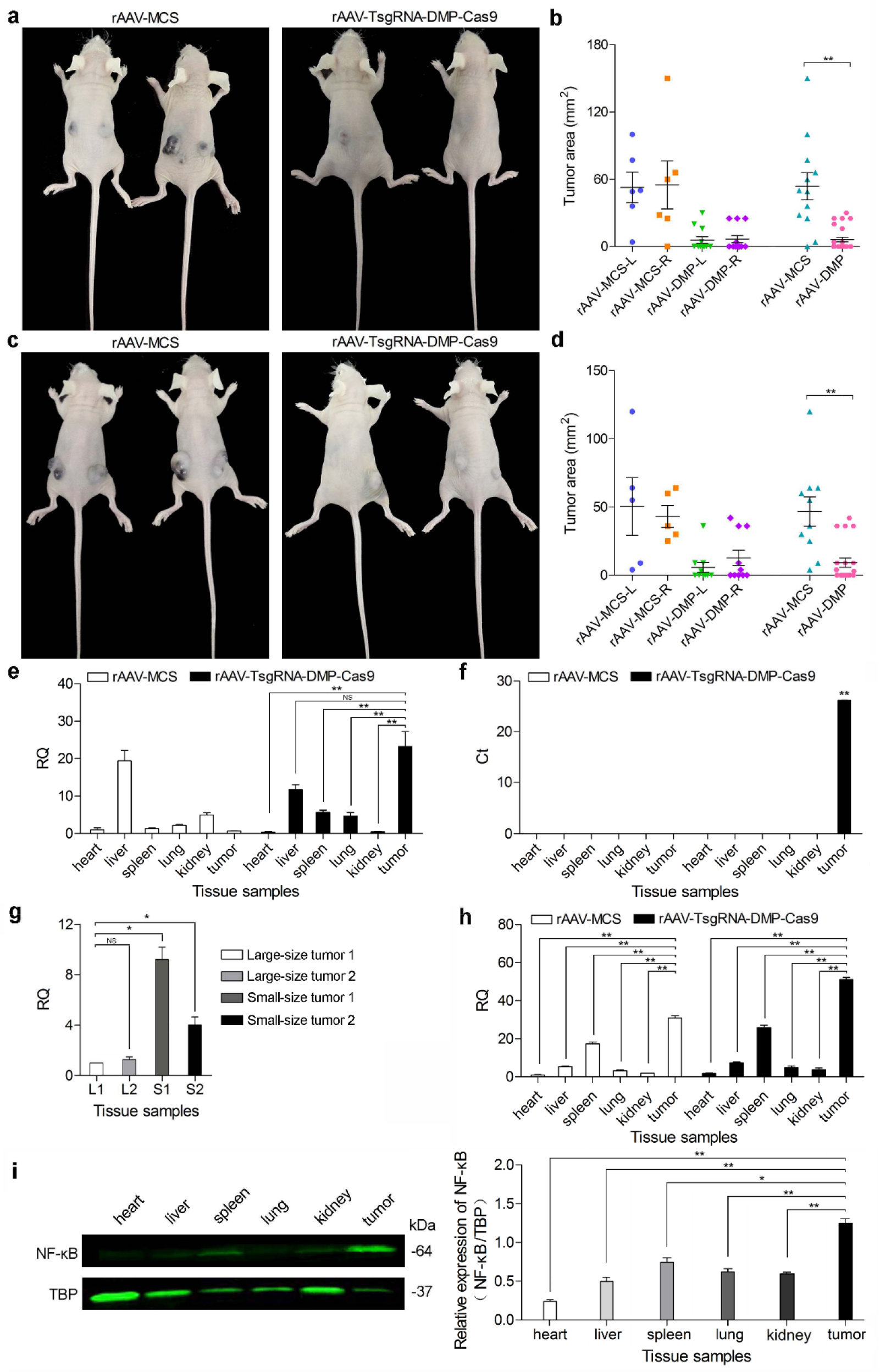
Gene therapy of xenografted cancers in mouse with the recombinant AAV packaged with the Nage vector expressing Cas9 and TsgRNA (rAAV-TsgRNA-DMP-Cas9). a, Representative images of nude mice subcutaneously transplanted with Hepa1-6 cells mixed with rAAV-MCS or rAAV-TsgRNA-DMP-Cas9, respectively (Experiment 1). b, Tumor size of nude mice in Experiment 1. L, left side; R, right side. c, Representative images of tumor-bearing nude mice intravenously injected with rAAV-MCS or rAAV-TsgRNA-DMP-Cas9 (Experiment 2). d, Tumor size of nude mice in Experiment 2. L, left side; R, right side. e, Viral DNA abundance in variant tissues of mice in Experiment 2. f, Relative quantity (RQ) of RelA mRNA in variant tissues of mice in Experiment 2. g, Ct values of qPCR detection of Cas9 mRNA in variant tissues of mice in Experiment 2. h, Viral DNA abundance in tumors with different sizes. Tumors were of mice Experiment 2. L, large-size tumor; S, small-size tumor. 1, tumor 1; 2, tumor 2. i, Western blot detection of abundance of NF-κB p65 protein in variant tissues. Tissues were samples form tumor-bearing mice. TBP was used as a loading control of nuclear protein to normalize the abundance of NF-κB. The quantified relative NF-κB protein abundance in variant tissues was shown beside the Western blot image. The statistical significance was checked between the tumor and other tissues. *, *p* ≤ 0.05; **, *p* ≤ 0.01.

In the second animal experiment, all mice were firstly subcutaneously transplanted with the Hepa1-6 cells on both sides of abdomen to produce tumor-bearing mice. After one week, the mice were randomly divided into two groups and intravenously injected with rAAV-MCS and rAAV-TsgRNA-DMP-Cas9, respectively. After one week, the mice were sacrificed for photographing and measuring the tumor size. The results revealed that all tumor-bearing mice injected with rAAV-MCS appeared large tumor on both sides (Figure 6c; Figure S5a). The mean tumor size of left and right sides was up to 202.2 mm^3^ and 130.3 mm^3^, respectively (Figure 6d). However, the tumor size of tumor-bearing mice injected with rAAV-TsgRNA-DMP-Cas9 was small in comparison with the control group. The mean tumor size of left and right side was only 14.85 mm^3^ and 39.95 mm^3^, respectively (Figure 6d). Some tumor were even eradicated (Figure 6c; Figure S5b). These results also demonstrated the strong tumor inhibition function of rAAV-TsgRNA-DMP-Cas9 by single intravenous administration. In addition, no mouse died in two animal experiments after rAAV administration, indicating the safety of rAAV vectors.

To further explore the cancer cell-specific expression of effector gene in the Nage vector, we collected the main tissues of mice in the second animal experiment, including heart, liver, spleen, lung, kidney, and tumor tissues. We detected the virus skeleton DNA and Cas9 mRNA in these tissues with qPCR. The results indicated that the viral skeleton DNA that represents virus distribution in tissues was in high abundance in the liver and tumor tissues (Figure 6e), indicating the tissue tropism of rAAV. There was also little distribution of rAAV in other tissues (Figure 6e). However, the qPCR detection of Cas9 expression demonstrated that Cas9 was only expressed in tumors (Figure 6f), indicating the tumor-specific expression of Nage vector. In order to preliminarily explore the factors leading to different tumor inhibition efficiency, two large and two small tumors were collected from the mice of the second animal experiments. The virus abundance in these tumors was detected with qPCR. The results indicated that there was more viruses in small tumors than in large tumors (Figure 6h). It suggested that the *in vivo* tumor inhibition efficiency is related to the abundance of AAV vectors in tumors.

Finally, to further explore the correlation between effector gene expression of Nage vector and NF-κB activity, we also detected the abundance of NF-κB RelA mRNA in variant tissues of mice of the second animal experiment by qPCR. The results indicated that NF-κB was transcribed at the highest level in tumors (Figure 6h). NF-κB was also transcribed at a higher level in spleen than in other tissues (Figure 6h). However, there was significant difference between tumor and spleen (Figure 6h), indicating high transcription of NF-κB in cancers. To further detect the activation of NF-κB in tissues, we prepared the nuclear extracts from heart, liver, spleen, lung, kidney, and tumor tissues of tumor-bearing mice. We then detected the NF-κB protein abundance in nuclear extracts with Western blot by using a anti-RelA antibody. The results indicated that a single 64-kDa band of NF-κB RelA protein was detected in liver, spleen, lung, kidney, and tumor tissues (Figure 6i). The relative NF-κB protein abundance in nuclear extracts of variant tissues was compared after normalized against TBP. The results indicated that there was the highest NF-κB activity in tumors (Figure 6i). Interestingly, it was found that there was also higher NF-κB activity in spleen than in other tissues (Figure 6i). However, there was significant difference between tumor and spleen (Figure 6h), indicating over activation of NF-κB in cancer. In combination with the result of Cas9 expression only in tumors, it can be concluded that only the NF-κB over activity in tumors can activate the expression of effector gene in Nage vector. Despite there was NF-κB activity in other tissues including liver, kidney, lung, and spleen, the effector gene expression was not activated by these relative low NF-κB activities, even not by the relatively high NF-κB activity in spleen. The results suggested that the effector gene expression of Nage vector is dependent on over activation of NF-κB in cancer, suggesting the *in vivo* safety of the Nage strategy.

## 4. Discussion

Contrary to the traditional strategy of inhibiting NF-κB activity, this study developed a novel strategy of utilizing NF-κB activity. In this strategy, an engineered NF-κB-specific promoter (DMP) was employed to drive gene expression. Activated by the constitutive NF-κB activity in cancer cells, an exogenous gene could be specifically expressed in cancer cells. To validate the strategy, we firstly demonstrated the cancer cell-specific over activation of NF-κB by detecting the NF-κB abundance in nuclear extracts of twelve cell lines. We then verified the strategy by expressing a reporter gene ZsGreen in twelve cell lines, which demonstrated that ZsGreen could only be specifically expressed in all detected cancer cell lines. We next verified the strategy by expressing CRISPR/Cas9 protein in twelve cell lines, which demonstrated that the Cas9 protein could also be specifically expressed in all detected cancer cell lines (reported by EGFP in Figure S1). Moreover, the Cas9 protein could induce death of all detected cancer cells by associating with a telomere-targeting sgRNA (TsgRNA). Because this strategy activated the expression of exogenous genes by depending on intracellular NF-κB activity, we thus named it as NF-κB-activating gene expression (Nage). In the Nage vector, the expression of a downstream effector gene was controlled by a NF-κB-specific promoter (DMP), which was constituted by a NF-κB decoy sequence and a minimal promoter sequence. Finally, we verified that the new strategy could be used to inhibit tumor growth in mice, in which a Nage vector co-expressing Cas9 protein under the control of DMP and TsgRNA under the control of U6 promoter was packaged in AVV viron.

To induce cancer cell death by using the Nage strategy, we used telomere DNA as target. The human telomere DNA consists of thousands of tandem repeats of TTAGGG [47]. Telomere functions in chromosome positioning, integrity and genomic stability through prohibiting nucleolytic degradation, chromosomal end-to-end fusion and irregular recombination [48]. In humans, the average length of telomere ranges from 10-15 kb [49]. However, the telomere DNA loses about 50-200 bp during per cell division due ot the inability of DNA polymerase to replicate the ends of liner DNA [50]. In general, the critically short telomere length can trigger cell death [51]. Therefore, telomere length acts as the mitotic clock for lifespan of eukaryotic cells [52]. However, due to the re-activation of telomerase, cancer cells can repair telomere DNA shortening, by which cancer cells become immortal [53]. Telomerase is a natural RNA-containing enzyme that can synthesize the repetitive telomere sequences [54], which thus helps to maintain the integrity of the genome in some cells such as embryonic stem cells [55]. Telomerase is silenced in most normal tissue cells but reactivated in most human cancer cells [56, 57]. Therefore, telomerase has been a attracting target to cancer therapy [58, 59]. Many telomerase inhibitors has thus been developed to treat cancers; however, none of them becomes the applicable clinical drugs due to their side effects [59–61]. On the other hand, a recent study indicated that the telomeric DNA damage could promote cancer cell death [62]. Therefore, we targeted telomere DNA to induce cancer cell death by Nage vector in this study.

CRISPR/Cas9 is a currently widely used gene editing tool to produce double-strand break (DSB) in genomic DNA [63–66]. In this study, we therefore used Cas9 as effector gene, which can introduce DSB to telomeric DNA by being guided by a telomere-targeting sgRNA. Our detection of telomere length indicated that the transfection of TsgRNA-DMP-Cas9-EGFP plasmid induced the shortening of telomere of transfected cancer cells; however, it produced no effect on telomere of normal cells. Because the TTAGGG repeats are the unique sequence of human telomere DNA, the genome blast showed that the TsgRNA target sequence is uniquely mapped to human telomere sequences. Therefore, the human telomere is the unique target of TsgRNA. The same TsgRNA has also been used to specifically image telomere with CRISPR/dCas9-EGFP [46]. Another reason to use telomere as target of Cas9 in this study is that human telomere DNA is long double-stranded DNA consisting of a multitude of tandem repeats of TTAGGG. Therefore, the human telomere could be multiply cut by Cas9-TsgRNA, which could rapidly produce enough DNA damage to promote cancer cell death. The previous studies revealed that TsgRNA could guide dCas9-EGFP to array of positions on telomere, by which telomere could be imaged with CRISPR/dCas9 [46]. Beside Cas9, many other genes that can induce cell apoptosis, differentiation, and suicide can also be used as effector genes in Nage vector.

To investigate if the Nage vector could be used to inhibit tumor growth *in vivo* by expressing TsgRNA and Cas9, we compacted the whole gene expression cassette including U6 promoter, TsgRNA, DMP, and Cas9 into AAV vector. We showed that the rAAV could efficiently inhibit tumor growth in model mice. AAV is now a widely used clinical gene therapy vector that has several significant advantages, including the tropism of both dividing and non-dividing cells, no host genome integration, stable long-term transgene expression, high production yields, and low immunogenicity [67]. In the past 2017, the US Food and Drug Administration (FDA) has approved several gene therapy such as hemophilia [68] and spinal muscular atrophy [69]. All of these gene therapy clinical tests use AAV as gene vector due to its long-standing safety [70]. However, a disadvantage of AAV vector is its limited packaging capacity (~4.7 kb). Despite long length of Cas9 protein coding sequence, due to short length of DMP, U6 promoter, and sgRNA coding sequence, we compacted the whole Nage vector cassette that expressing TsgRNA and Cas9 into a AAV viron. This single AAV viron package of full-length Nage vector is beneficial for increasing virus administration dosage and enhancing therapeutic effect. This is better than the current multiple AAV combinant *in vivo* application due to large size of CRISPR/Cas9 system and limited package capacity of AAV vector [71]. Because DMP is a weak promoter, the capability of stable long-term transgene expression capability of AAV vector is helpful for the sufficient expression of Cas9 proteins. Also due to the long-term transgene expression capability of AAV vector, as performed in the current clinical gene therapies, we performed single administration of rAAV to cancer therapy of tumor-bearing mice. The serotype of AAV used in this study is AAV2, which is an liver-tropism AAV. In this study, we transfected human hepatoma HepG2 and mouse hepatoma Hepa1-6 cells *in vitro* with rAAV. We treated the mice bearing the tumors caused by Hepa1-6 inoculation. It is worthy to mention that no mice died when two animal experiments, indicating safety of rAAV.

In this study, we also used the 293T cell in detecting NF-κB activity, transfection DMP-ZsGreen and TsgRNA-DMP-Cas9-EGFP. We found that there were the highest NF-κB activity (Figure 1) and expression of ZsGreen (Figure 2) and EGFP (Figure S2) in this cell. However, the transfection of TsgRNA-DMP-Cas9-EGFP did not induce the significant death of the cell (Figure 3). The detection of telomere length revealed that the transfection of TsgRNA-DMP-Cas9-EGFP also did not shorten the telomere length of the cell (Figure 4). This is an unexpected but interesting result. It is known that HEK293 cell line was derived from the normal human embryonic kidney tissue cultured *in vitro*. 293T cell line was derived from HEK293 cells after a series of transfection with sheared gDNA of Adenovirus 5 [72, 73]. The “T” means that it expresses the Large T antigen which is important for replicating plasmids containing an SV40 origin of replication to high copy number in the transfected cell. These facts suggested that 293T cell is not a normal cell line as MRC5 and HL7702. However, despite its abnormal chromosome number (64) and structure, it has also never been used as a cancer cell line. Therefore, the underlying reason why TsgRNA-DMP-Cas9-EGPF transfection did not induce death of this cell should be further explored in the future.

In this study, we found that there was also basic NF-κB activity in normal organs, including liver, lung, kidney, and spleen. This may be reason why the traditional NF-κB inhibitors have unacceptable side effects. Despite the basic NF-κB activity in these normal organs, we found that the effector gene expression of Nage vector was not activated by these relative low NF-κB activities, especially not by the relatively high NF-κB activity in spleen. The results showed the high cancer specificity of effector gene expression of Nage vector. The relative high NF-κB activity in spleen should be related with the major physiological function of NF-κB in immunity. Spleen is the biggest immunity. There is a large number of lymphocytes and macrophage in spleen. As we discussed above, DMP is a weak promoter. According to the results of this study, DMP can not be activated by the relative low NF-κB activities in normal organs. This is critical for the clinical application of Nage vector as a gene therapy of cancers.

In summary, we developed a novel gene therapy of cancer based on a newly discovered cancer cell-specific gene expression technique, Nage. This technique fully utilizes the widely over-activated NF-κB activity in cancer cells, which is completely opposite to the current strategy that inhibits NF-κB activity with various inhibitors. Until now, there is still no Nage-like strategy has been developed. Therefore, this is the first time to develop a NF-κB utilizing strategy for cancer therapy, which may be also used to treat other NF-κB over activated diseases such as inflammation [12]. By changing effector gene in Nage vector, variant gene therapies can be developed to treat the NF-κB over activated diseases.

## Author contributions

J.K.W. conceived the study and designed experiments. W.D. designed and performed main experiments. J.W. prepared reagents and performed partial experiments. J.K.W. and W.D. wrote the manuscript with support from all authors.

## Acknowledgments

This work was supported by the National Natural Science Foundation of China (61571119) and the National Key Research and Development Program of China (2017YFA0205502).

## Supplementary information

**Table S1.**
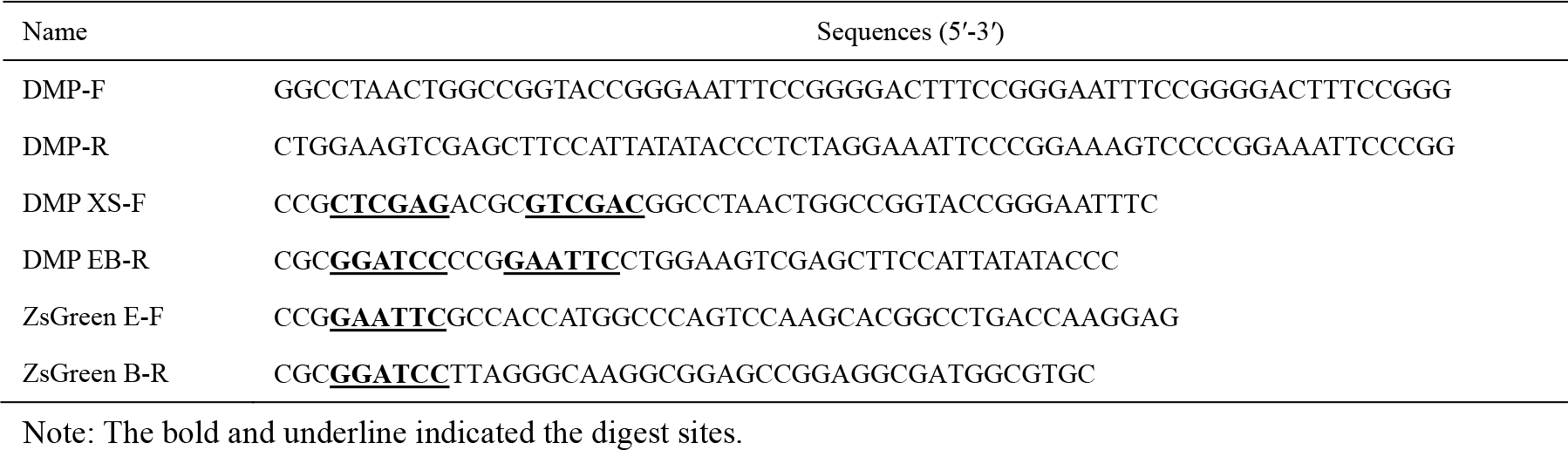
Oligonucleotides used in DMP-ZsGreen experiment.

**Table S2.**
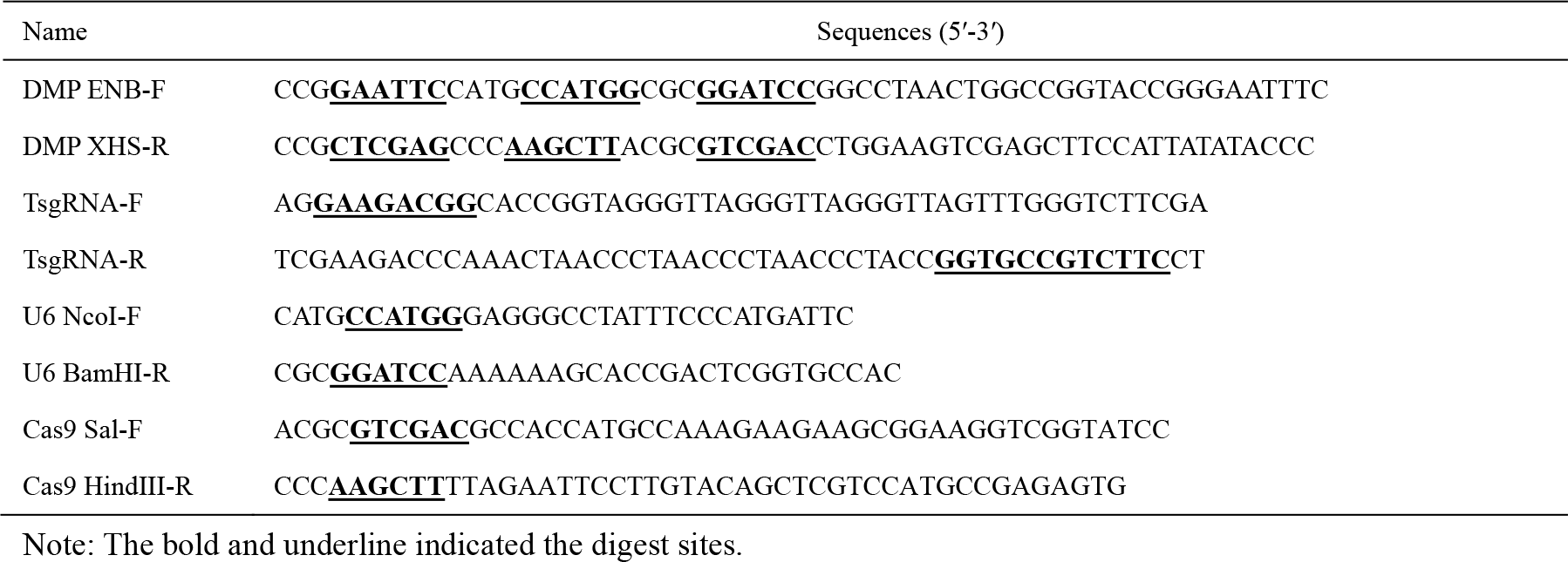
Oligonucleotides used in TsgRNA-DMP-Cas9-EGFP experiment.

**Table S3.**
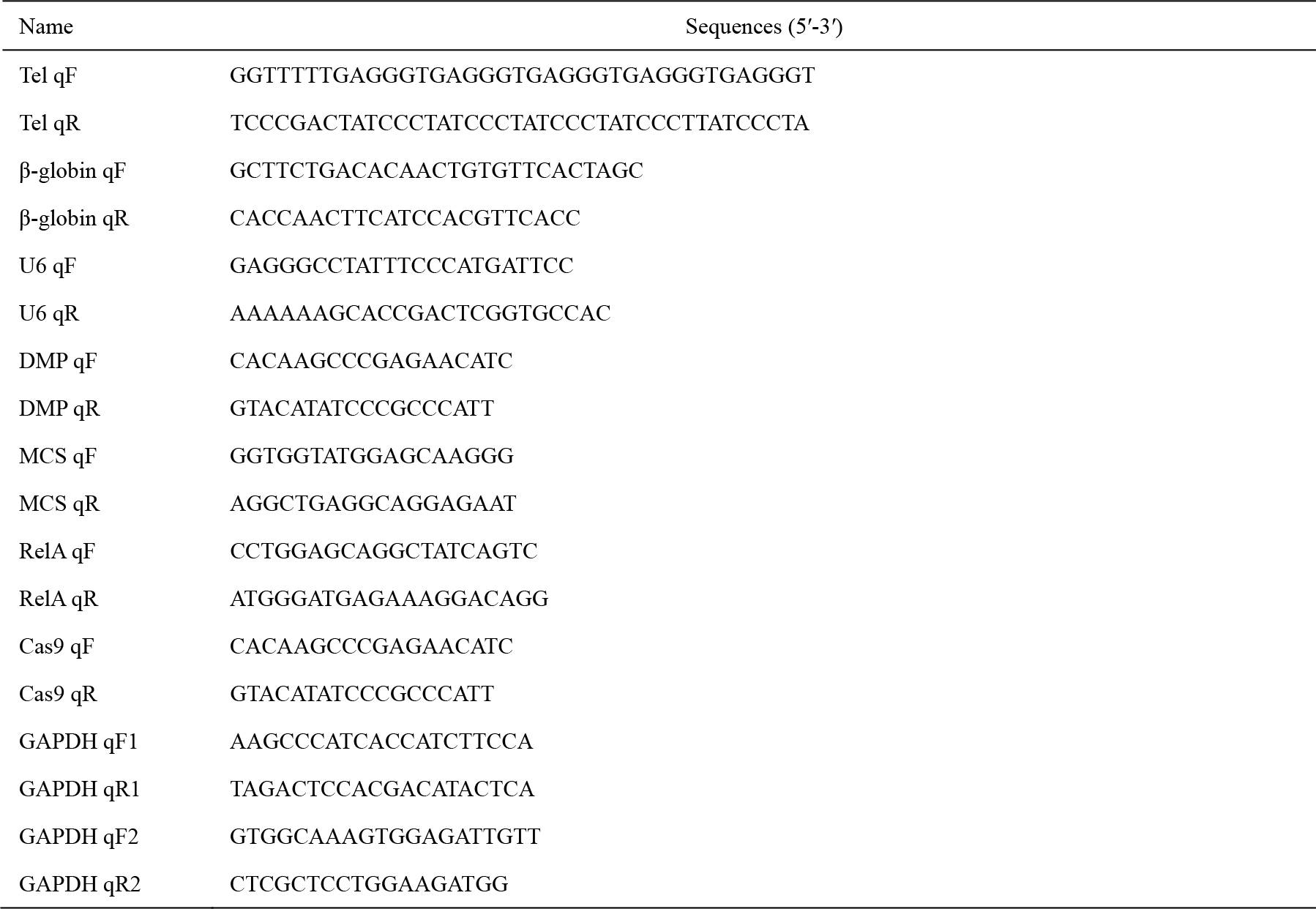
Oligonucleotides used in qPCR experiment.

**Figure S1.**
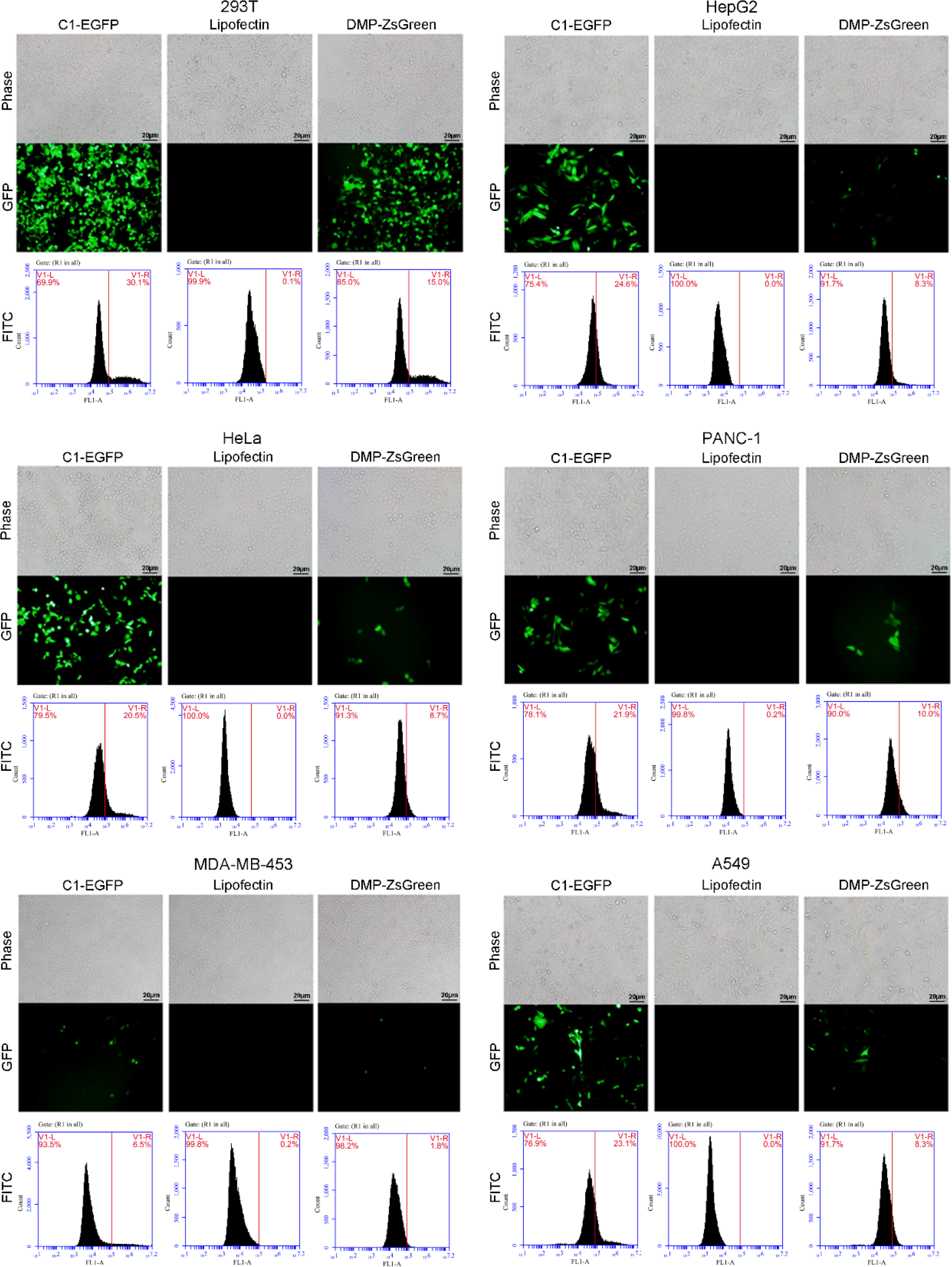
Evaluation of the cancer cell specificity of Nage vector. Representative cell images under the bright field (Phase) and fluorescence (GFP). Cells were transfected by pC1-EGFP and pDMP-ZsGreen, respectively. In pC1-EGFP, the expression of EGFP is controlled by a strong promoter CMV. The pC1-EGFP transfection was used to monitor the transfection process as a positive control. Cells transfected by lipofectin were used as blank control. The results of DMP-ZsGreen were also included in this figure for easily comparing the results of different treatments.

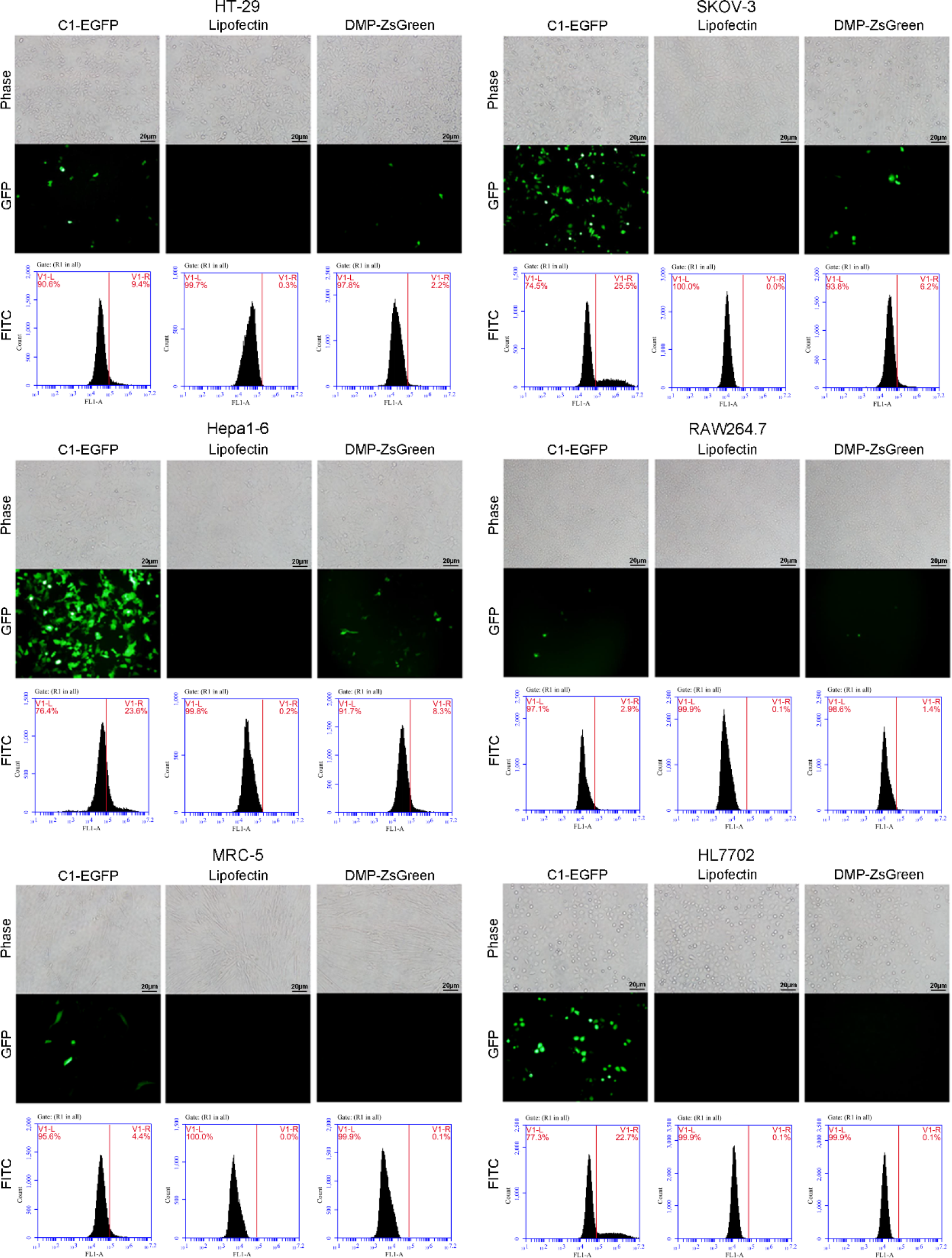

**Figure S2.**
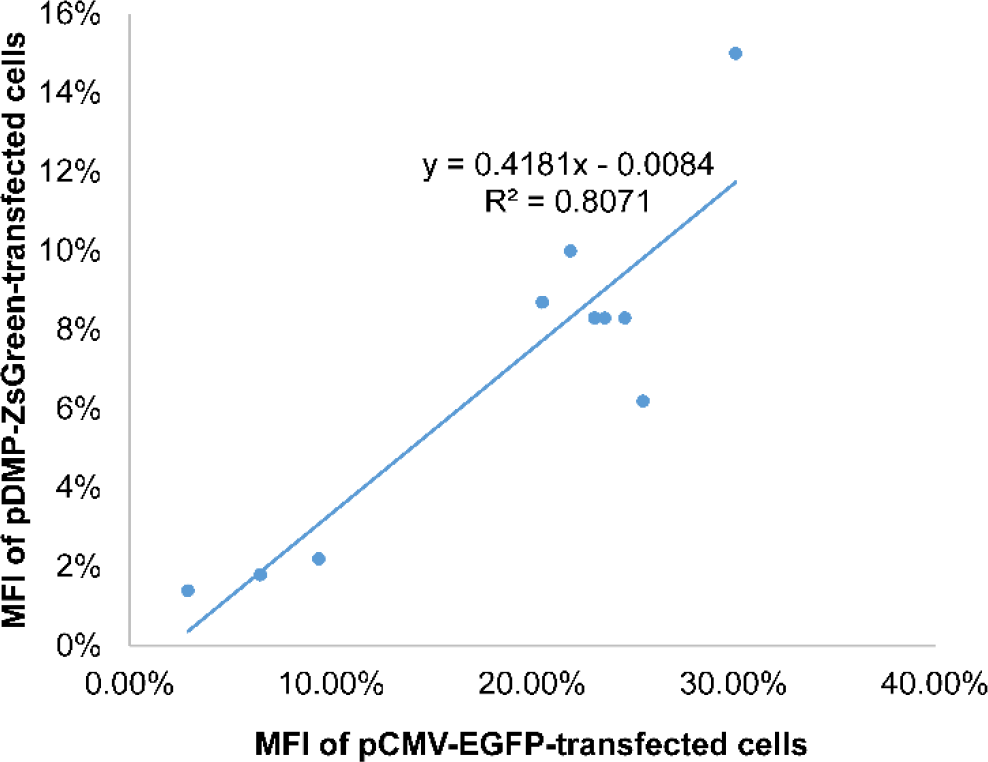
Correlation between the MFI of cells transfected by pDMP-ZsGreen and pCMV-EGFP (). The MFI data were shown in Figure S1.

**Figure S3.**
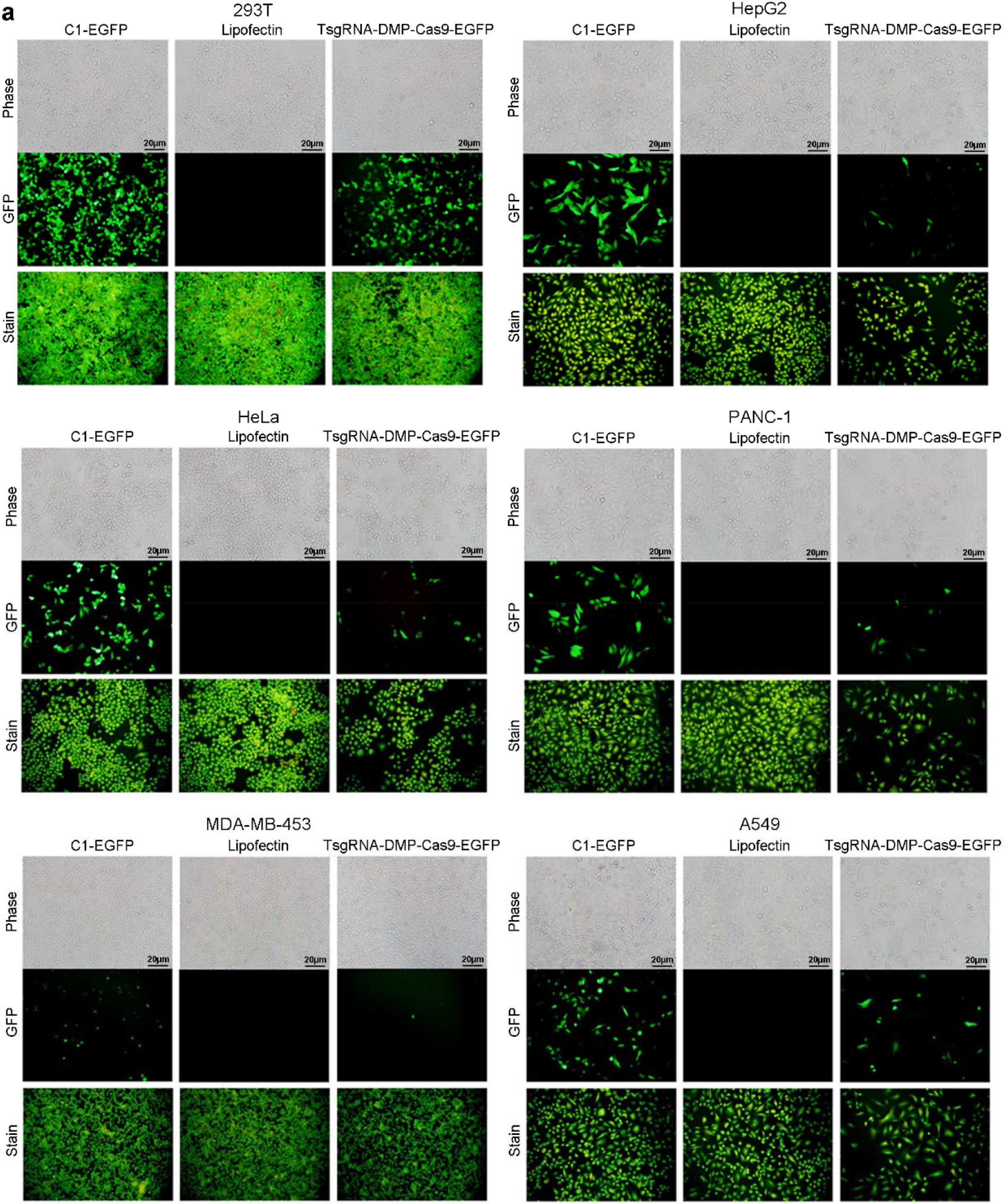
Killing cancer cells with Nage vector. a, Representative fluorescent images of cells. Cells were transfected by plasmid Nage vector TsgRNA-DMP-Cas9-EGFP and pC1-EGFP. Cells were imaged at the GFP channel for detecting EGFP expression. Cells were imaged at bright field for showing growth. For more clearly showing the cell growth, cells were then stained with acridine orange and imaged under fluorescence. b, Alamar blue assay of cell viability. The pC1-EGFP transfection was used to monitor the transfection process as a positive control. Here, the pC1-EGFP transfection was also used as a negative control to show the effect of TsgRNA-DMP-Cas9-EGFP transfection on cell growth. Cells only transfected by lipofectin were used as blank control. The results of TsgRNA-DMP-Cas9-EGFP transfection were also included in this figure for easily comparing the results of different treatments.

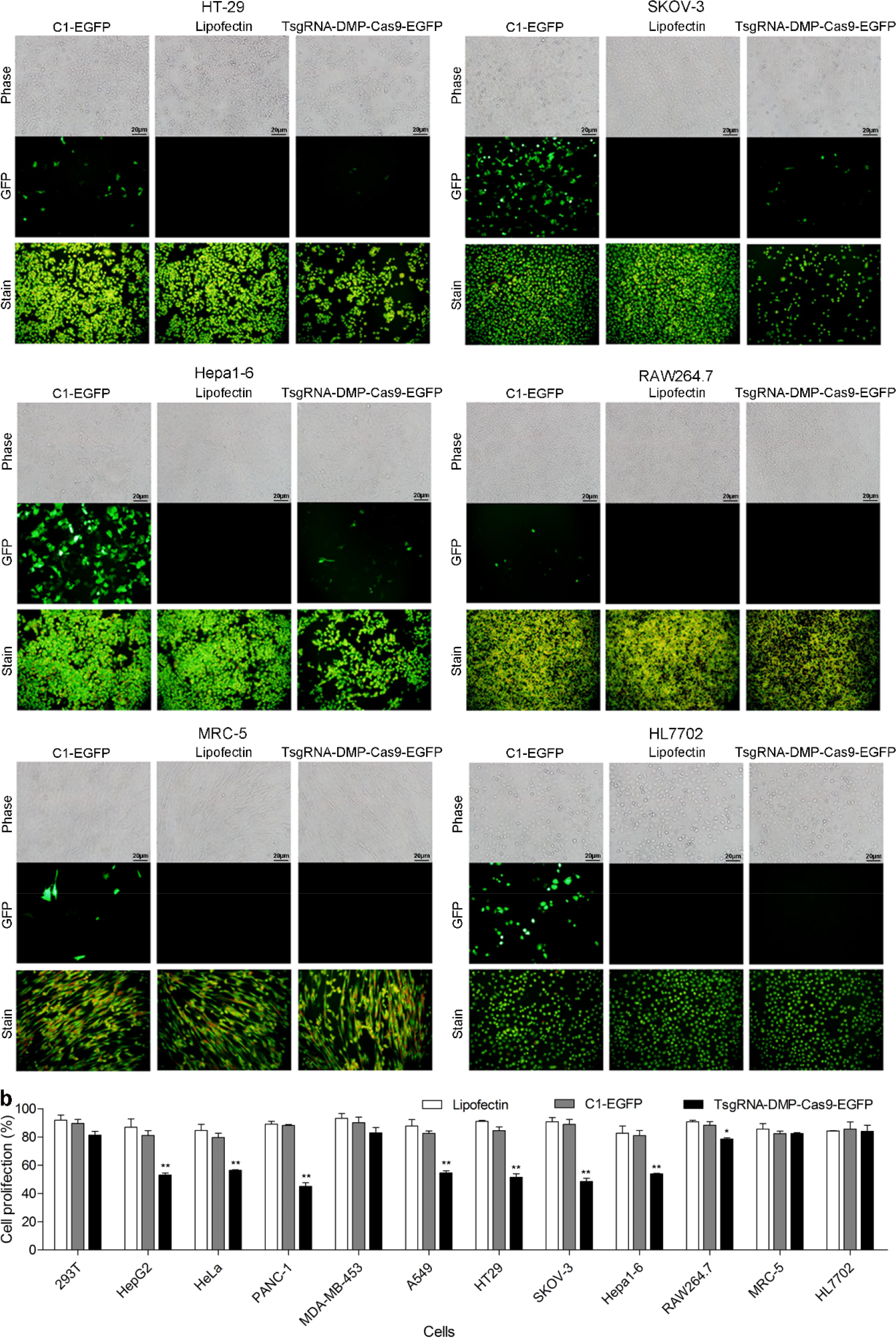

**Figure S4.**
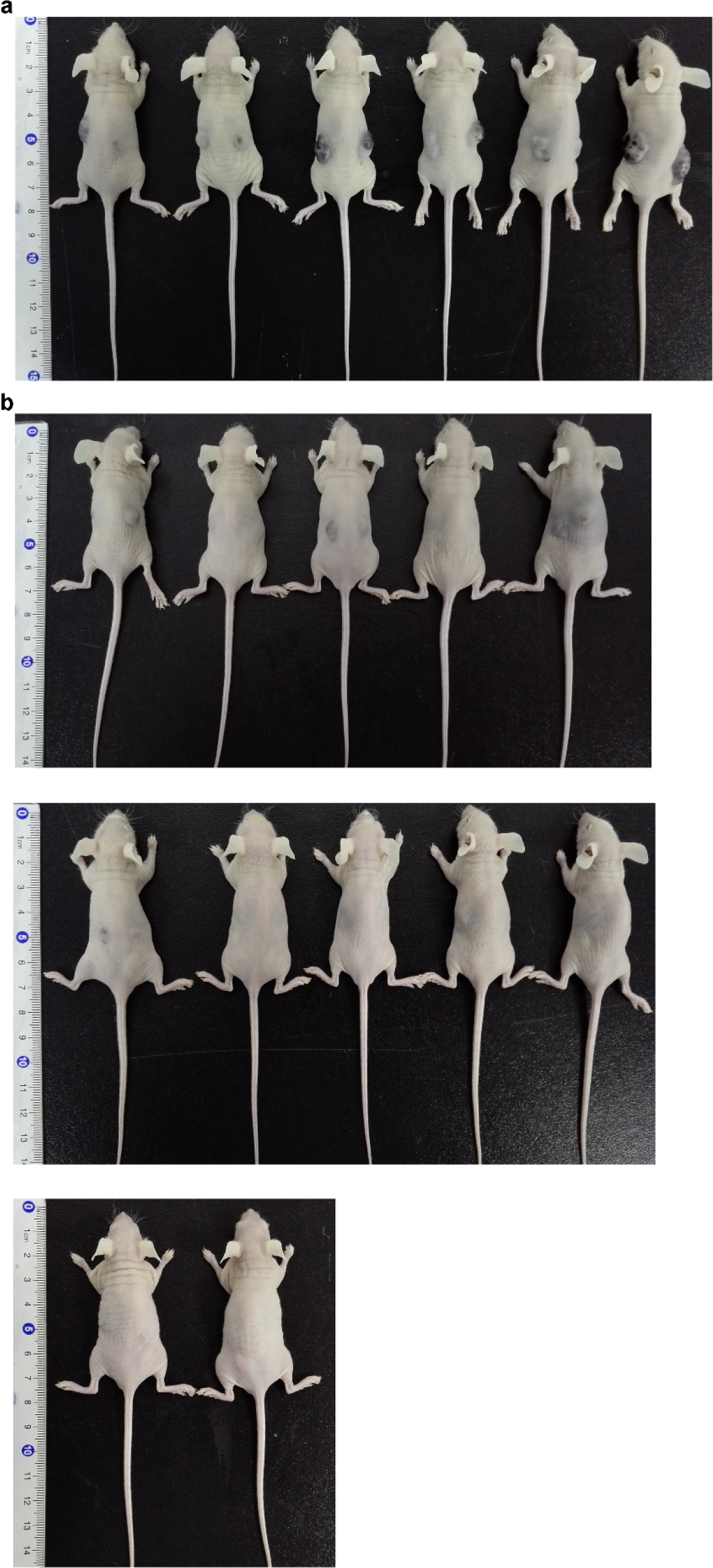
Application of Nage vector for cancer therapy by transplanting the mixture of Hepa1-6 cells and rAAV. a, Nude mice subcutaneously transplanted with the mixture of Hepa1-6 cells and rAAV-MCS. b, Nude mice subcutaneously transplanted with the mixture of Hepa1-6 cells and rAAV-TsgRNA-DMP-Cas9.

**Figure S5.**
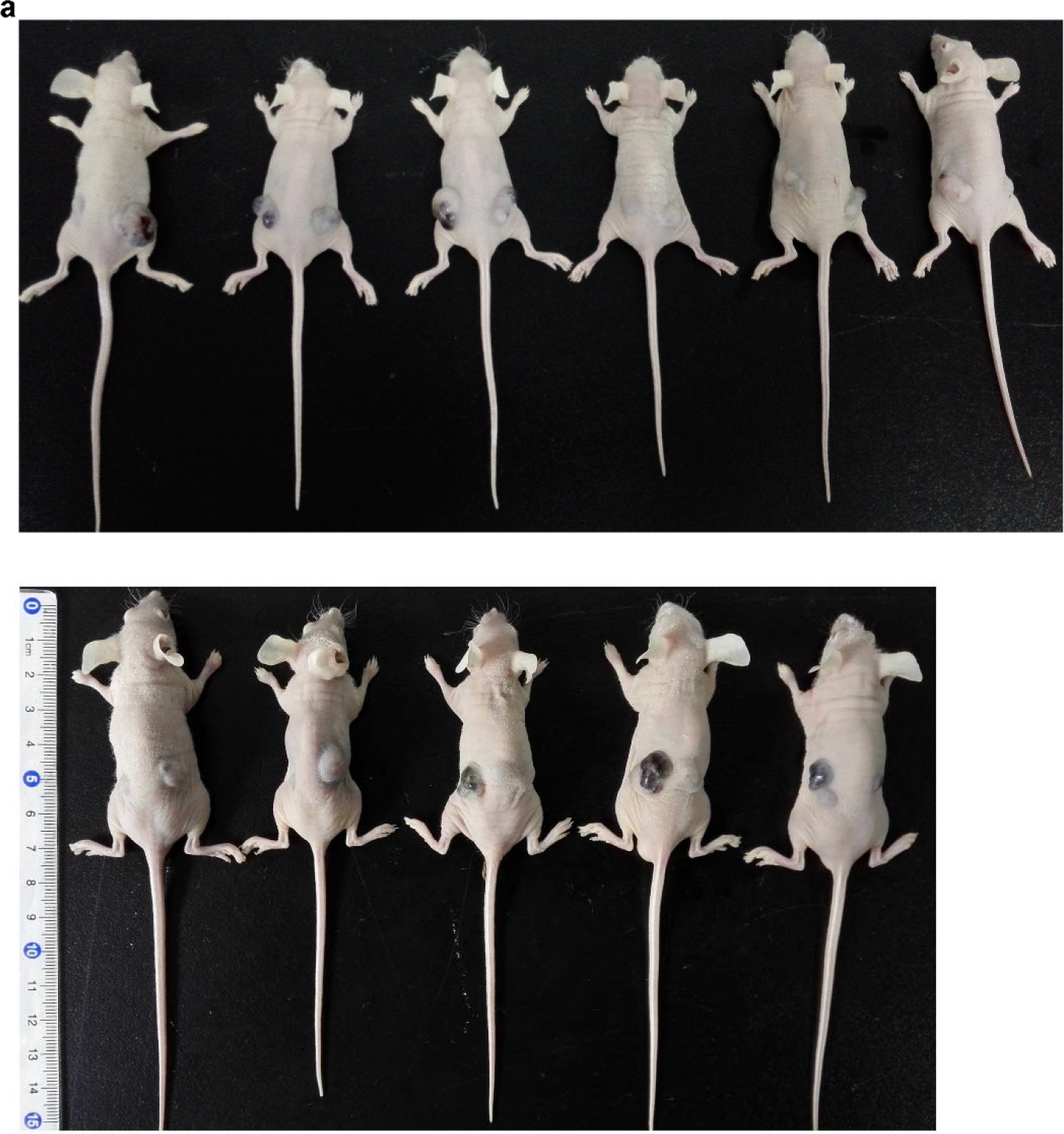
Application of Nage for cancer therapy by intravenously injecting rAAV into tumor-bearing mice. a, Tumor-bearing nude mice intravenously injected with rAAV-MCS. b, Tumor-bearing nude mice intravenously injected with rAAV-TsgRNA-DMP-Cas9. The tumor-bearing nude mice were prepared by subcutaneously transplanting with Hepa1-6 cells.

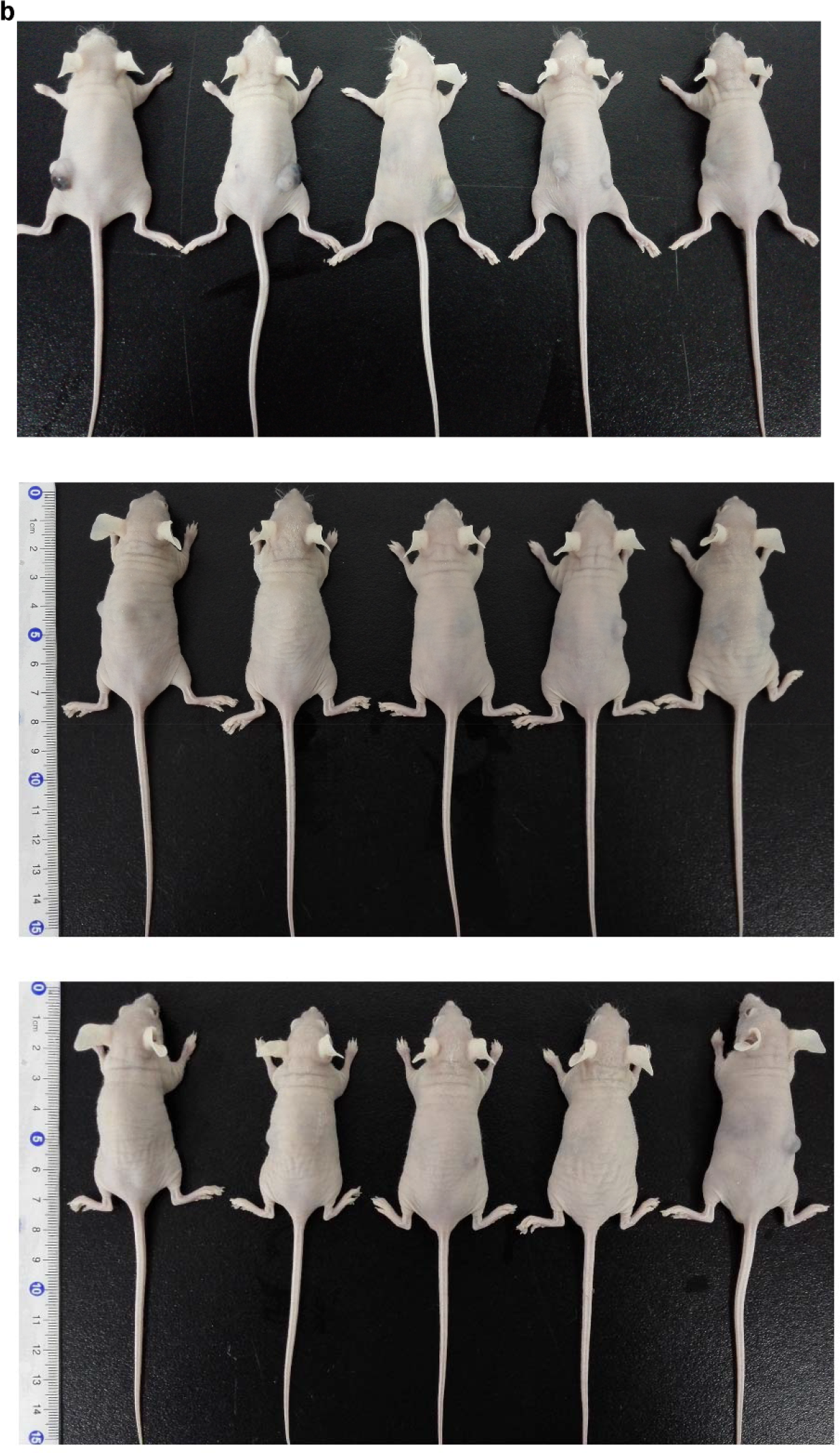

### Supplementary Sequences

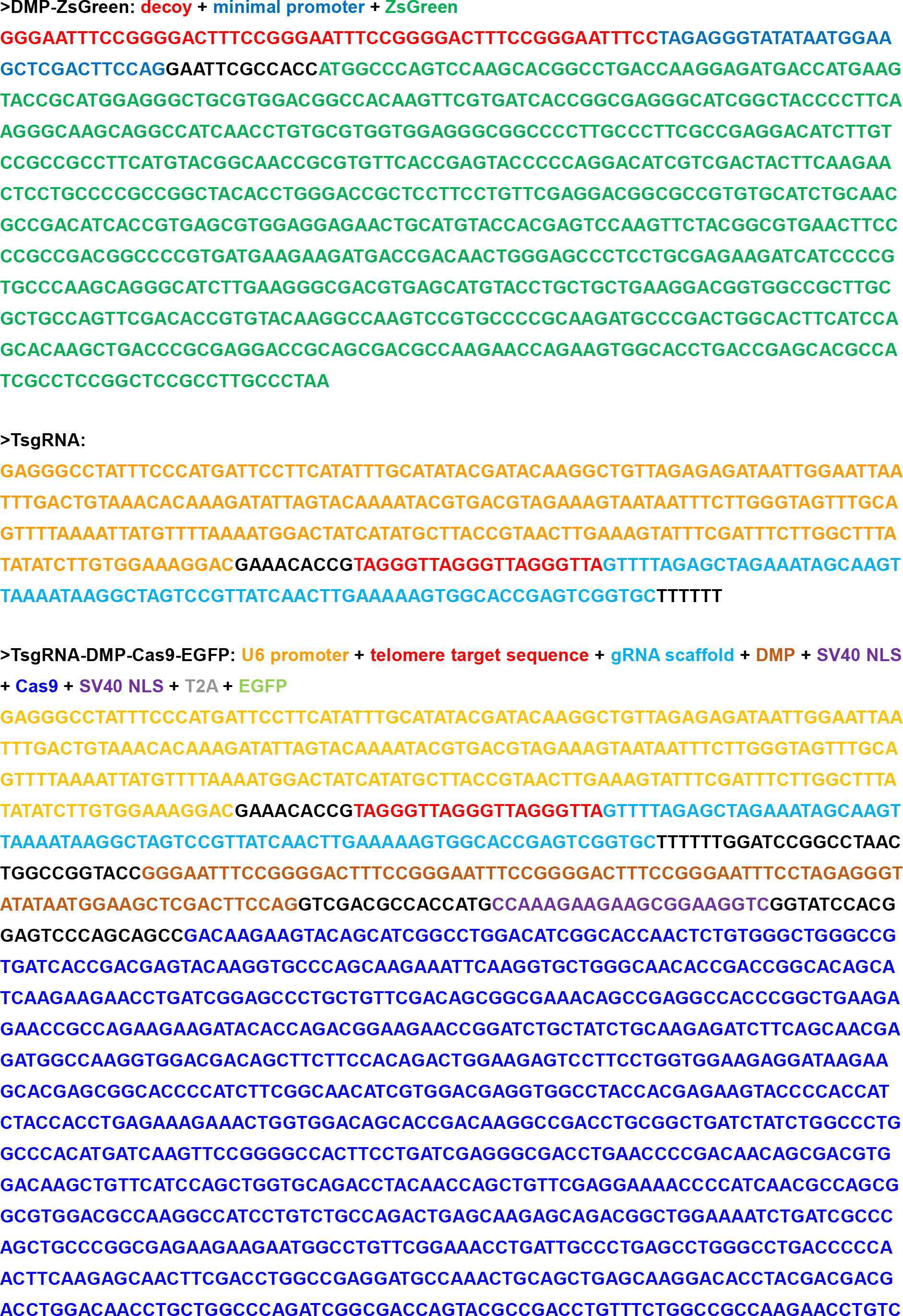

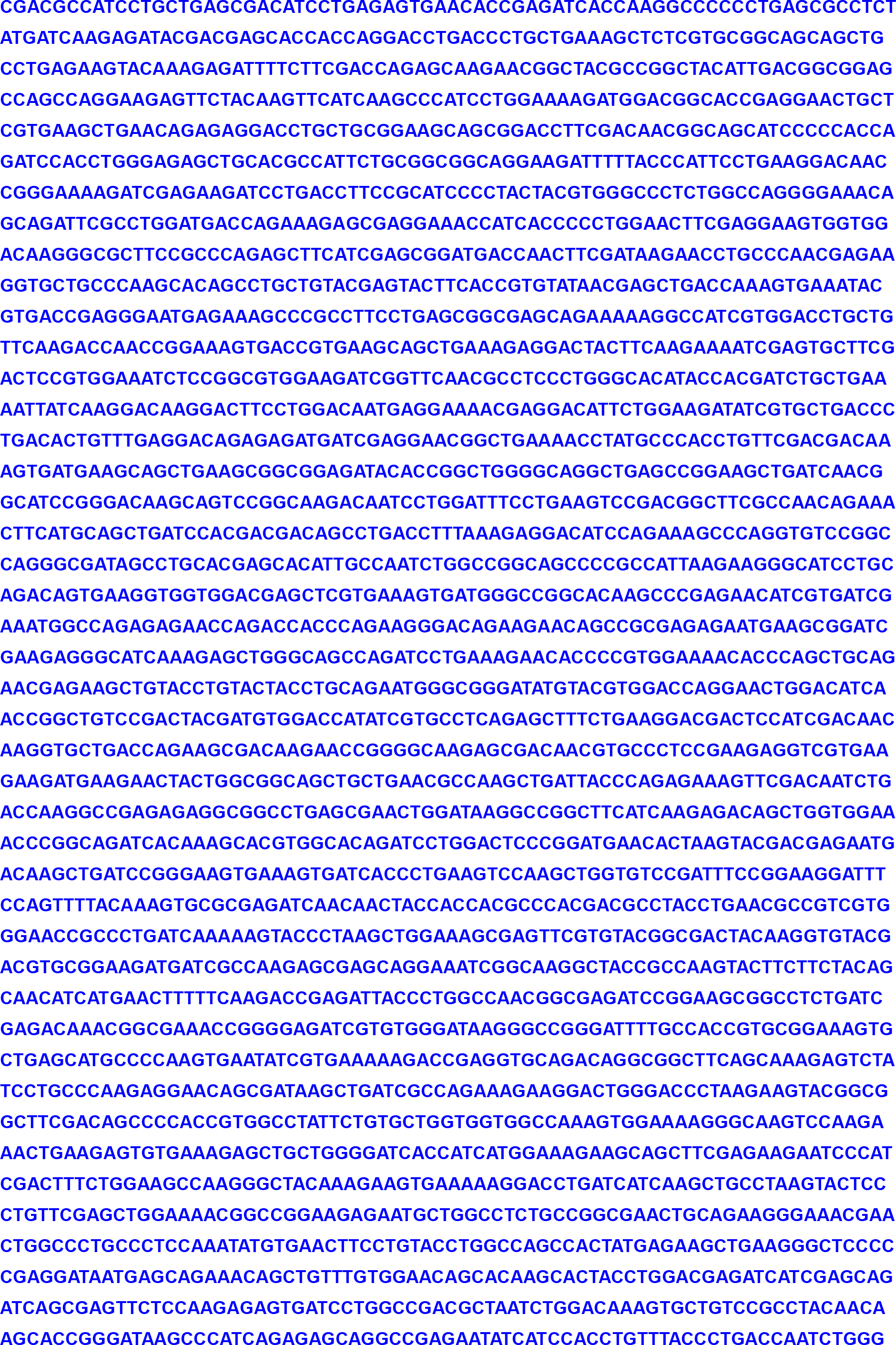

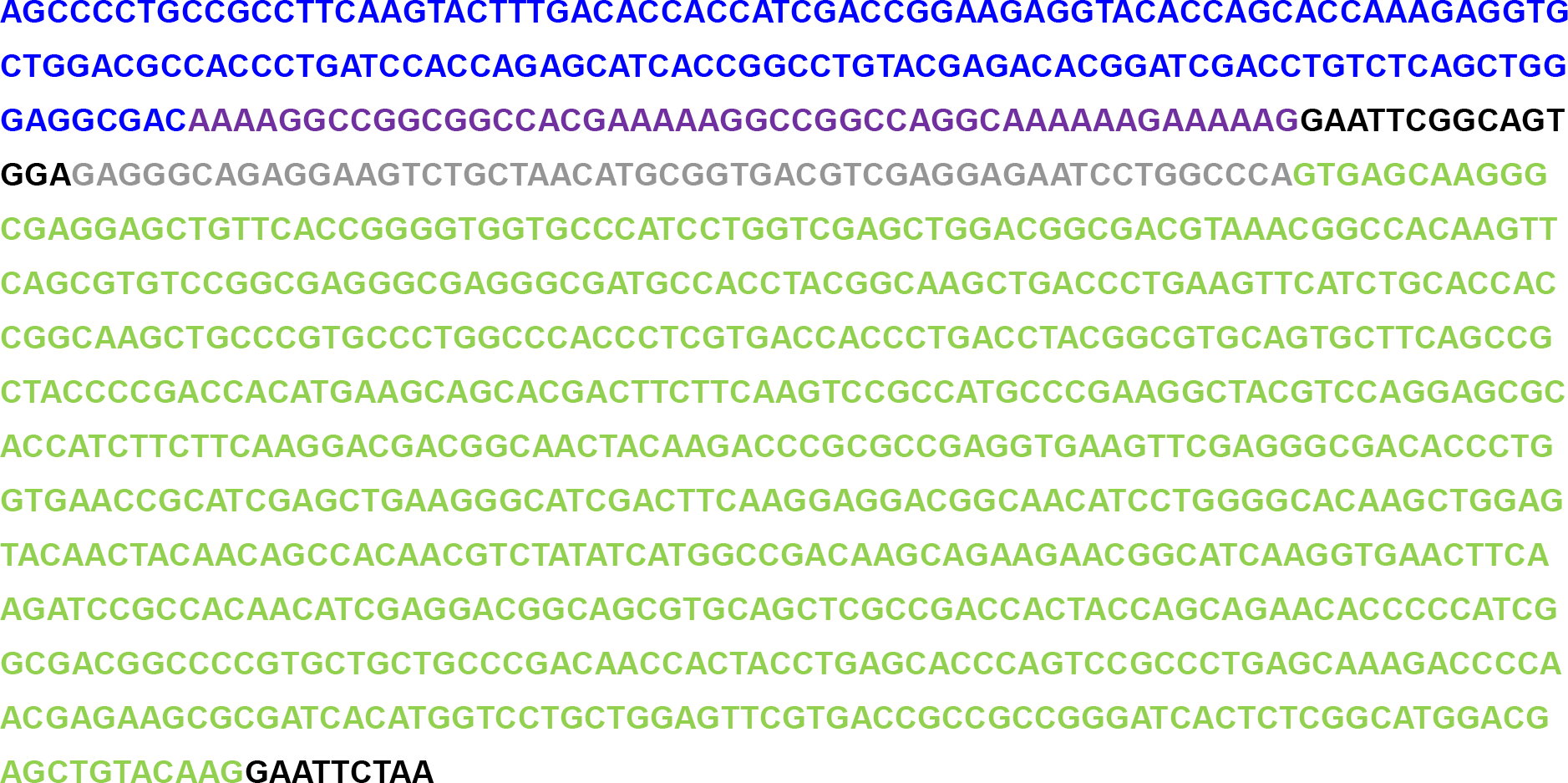

